# Anatomical and behavioral correlates of auditory perception in developmental dyslexia

**DOI:** 10.1101/2023.05.09.539936

**Authors:** Ting Qi, Maria Luisa Mandelli, Christa L. Watson Pereira, Emma Wellman, Rian Bogley, Abigail E. Licata, Edward F. Chang, Yulia Oganian, Maria Luisa Gorno-Tempini

**Author notes:** Correspondence to: Ting Qi and Yulia Oganian, Full address: Department of Brain Cognition and Intelligent Medicine, Beijing University of Posts and Telecommunications, Beijing, China; Center for Integrative Neuroscience, University of Tübingen, Tübingen, Germany. Shared senior authorship (Yulia Oganian, Maria Luisa Gorno-Tempini).

## Abstract

Developmental dyslexia is typically associated with difficulties in basic auditory processing and in manipulating speech sounds. However, the neuroanatomical correlates of auditory difficulties in developmental dyslexia (DD) and their contribution to individual clinical phenotypes are still unknown. Recent intracranial electrocorticography findings associated processing of sound amplitude rises and speech sounds with posterior and middle superior temporal gyrus (STG), respectively. We hypothesize that regional STG anatomy will relate to specific auditory abilities in DD, and that auditory processing abilities will relate to behavioral difficulties with speech and reading. One hundred and ten children (78 DD, 32 typically developing, age 7-15 years) completed amplitude rise time and speech in noise discrimination tasks. They also underwent a battery of cognitive tests. Anatomical MRI scans were used to identify regions in which local cortical gyrification complexity correlated with auditory behavior. Behaviorally, amplitude rise time but not speech in noise performance was impaired in DD. Neurally, amplitude rise time and speech in noise performance correlated with gyrification in posterior and middle STG, respectively. Furthermore, amplitude rise time significantly contributed to reading impairments in DD, while speech in noise only explained variance in phonological awareness. Finally, amplitude rise time and speech in noise performance were not correlated, and each task was correlated with distinct neuropsychological measures, emphasizing their unique contributions to DD. Overall, we provide a direct link between the neurodevelopment of the left STG and individual variability in auditory processing abilities in neurotypical and dyslexic populations.

## Introduction

Developmental dyslexia (DD) is characterized by difficulties with reading and spelling that persist throughout life and cannot be attributed to general cognitive abilities or poor educational opportunities.^1^ While primarily diagnosed through reading performance, DD often involves deficits in phonological awareness, that is the ability to process and manipulate speech sounds.^2^ In this view, reading, which requires mapping from orthography to speech sounds (phonology), breaks down because of impaired speech sound representations or access to these representations (Phonological deficit theory of dyslexia).^3–5^

Speech-related deficits in DD are not exclusively phonological, and some theories frame them as stemming from a more general auditory processing deficit.^2,6,7^ In this view, general auditory impairments drive the inability to develop and access phoneme representations.^8,9^ This is in line with the view that to extract speech sounds from auditory streams, the auditory system has to identify a range of complex acoustic cues in the speech signal.^10^ A key acoustic feature for speech comprehension is amplitude modulations, specifically amplitude rises, which cue speech structure at phrasal and syllabic levels.^11^ Indeed, a large body of work has found impaired processing of amplitude rises in DD.^12–18^ Furthermore, individuals with DD sometimes show deficits in the perception of speech in noisy backgrounds, which is more challenging than under optimal listening conditions and might require more precise phoneme representations.^19,20^ Amplitude rise time deficits are also evident in infants at risk for dyslexia, and the predictive role of rise-time abilities for later vocabulary and phonological awareness is well established from an early age in typically developing children as well.^21,22^ In contrast, speech-in-noise deficits are not present in infants at familial risk for DD, but rather emerge during preschool and are found to improve substantially with age, continuing into late childhood in typically developing children.^23,24^ However, it remains debated whether these auditory deficits characterize all or only subgroups of individuals with DD and how they relate to each other, and to reading and phonological abilities.

These behavioral deficits are complemented by reports of atypical neuroanatomical patterns in auditory pathways in DD (see ^6,25–29^ for similar findings in the visual pathways).^3,30–33^ Among others, altered cortical thickness, myelinated cortical thickness ratio, and surface area lateralization of auditory temporal cortices have been reported in DD and in individuals with familial risk for DD.^34–40^ Notably, high variation is evident with respect to the specific locations and patterns found across studies. Specifically, a decrease in cortical thickness has been found in the anterior portion of the superior temporal gyrus (STG), whereas increases in cortical thickness have been reported in the right STG, middle temporal gyrus (MTG), and Heschl’s gyrus.^38,41^ Similarly inconsistent patterns were also observed for gyrification patterns.^42–45^ This range of observations supports a core role for the temporal cortex in DD, but also fuels the idea that the heterogeneity of behavioral deficits in DD may be mapped to variations in anatomical abnormalities. These anatomical changes might contribute to auditory impairments, with early sensory differences also playing a role in these variations.^46,47^ Complementing this, studies in typically developing children showed that the neuroanatomy of the left superior temporal cortex is crucial for reading, with better reading performance associated with greater gray matter volume and surface area and thicker cortical thickness due to its role in auditory processing of speech.^48,49^ Furthermore, auditory capacities are linked to the neuroanatomy of the temporal cortex, reinforcing its fundamental role in auditory perception.^50–53^

Overall, it remains unclear how neuroanatomical cortical structure, particularly in auditory temporal cortical areas, relates to specific auditory behavioral deficits in DD. Until recently, this gap was widened by our limited understanding of the neural computations underlying the processing of speech sounds in human auditory cortices. However, recent advances in intracranial electrophysiology (iEEG) recordings from auditory and speech cortices have revealed the rapid dynamics of cortical speech sound representations.^54^ Most relevant to the behavioral deficits in DD, recent studies established a spatial map for the encoding of amplitude rises at phrasal onsets in posterior STG (pSTG), and phonemes and syllabic amplitude rises in middle STG (mSTG).^11,55,56^ This detailed spatial brain map for speech sound processing opens new avenues for mapping auditory processing deficits in DD to underlying neural substrates. Indeed, non-invasive electrophysiology studies of DD found reduced neural responses to amplitude modulations in speech and non-speech sounds and some functional MRI studies report atypical activation of left hemispheric temporal regions in DD for a wide variety of stimuli and perceptual discrimination tasks.^3,57–61^

Here, we built on these findings to hypothesize that the ability to process and manipulate speech and non-speech sounds in developing populations depends on the neuroanatomical structure and development of the STG. To test this, we behaviorally assessed the ability to discriminate amplitude modulations in sounds and to perceive speech in noise, alongside cognitive, reading, and phonological abilities, in a group of children with a diagnosis of DD and in age-matched typically developing (TD) children. To test how these two auditory tasks map onto the brain’s structure we used anatomical MRI scans in the same cohort. We calculated the local gyrification index (LGI), which has been found to be the best cortical geometric measure discriminating between DD and neurotypical groups.^43^ Based on our prior intracranial results, we hypothesized that amplitude modulation processing abilities would be correlated with neuroanatomical structure in the pSTG, whereas we expected speech in noise perception to be associated with the mSTG. Further, we hypothesized that the abilities to process speech and non-speech sounds might be independent in DD.

## Materials and methods

### Participants

This study includes 78 children with DD and 32 TD children who successfully completed at least one of the auditory tasks. A subgroup of 102 (76 DD, 26 TD) completed the MRI session. To maximize sample sizes, each of the following analyses included the maximal subset of children that completed the relevant tasks (see Supplementary Table 1 for initial sample sizes and Table 1 for included sample sizes). All DD children were selected from the recruitment base at the UCSF Dyslexia Center, a multidisciplinary research program that performs neurological, psychiatric, cognitive, linguistic, and neuroimaging evaluations of children with language-based neurodevelopmental disorders. Of note, the center partners with several schools for young individuals with language-based learning differences, where children routinely participate in Orton-Gillingham-based intervention programs, characterized by highly structured training focusing on phonological awareness, phonics, fluency, vocabulary, and comprehension. TD children were recruited through local schools and parent networks. Reading and language abilities were assessed using a battery of standardized reading tests. General cognitive abilities were assessed using the Matrix reasoning test (WASI).^62^ All DD children were native speakers of English, aged between 7 and 15 years, and underwent a detailed clinical interview and neurological examination. The criteria for inclusion of DD were that a child had prior formal diagnoses of DD and, despite participation in extensive school-based reading intervention, currently at least one reading score falling below the 25^th^ percentile of same-aged peers on a standardized reading test and general cognitive abilities within the normal range (16^th^ percentile) of same-aged peers. Exclusion criteria for both groups included acquired brain injury, neurological disorders such as perinatal injuries, seizures, and severe migraine. TD were excluded if a single score on any of the reading or general cognitive ability tests fell below the normal range (16^th^ percentile) of same-aged peers (see Table 1). Further exclusion criteria for the TD group were a history of academic difficulties, prior diagnoses of DD, or other developmental, neurological, or psychiatric disorders. Behavioral assessments were typically completed within 6 months from MRI scans, with a *mean* interval of 0.09 (*SD* = 0.15, in years). Written informed consent was obtained from the legal guardian or parent of the children. Additionally, children provided verbal consent for participation before the experiments. The study was approved by the UCSF Committee on Human Research and complied with the declaration of Helsinki.

**Table 1.**
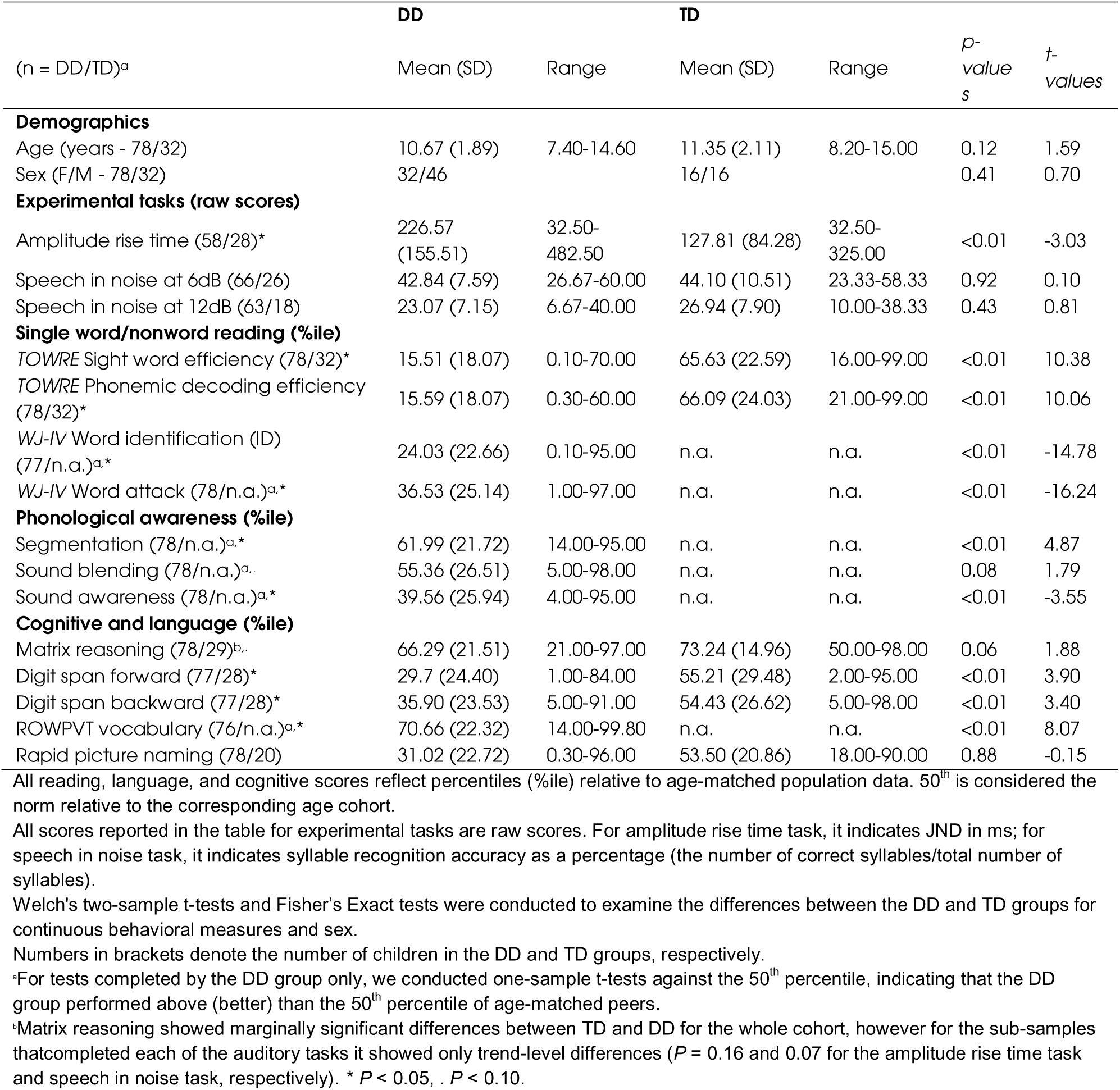
Demographics and behavioral characteristics of the developmental dyslexic (DD) and typically developing (TD) children.

### Neuropsychological and academic assessment

Children with DD underwent a comprehensive battery of neuropsychological and academic testing. Neuropsychological testing consisted of matrix reasoning (MR) for general cognitive abilities (WASI Matrix Reasoning)^62^, digit span forward (DSF) and backward (DSB) (WISC-IV Integrated Digit Span)^63^ for verbal short-term memory and working memory respectively, receptive one-word picture vocabulary Test-4 for general vocabulary skills (ROWPVT)^64^, and rapid picture naming for lexical processing (retrieval) speed (Woodcock-Johnson IV)^65^.

To evaluate their reading abilities, all DD children completed two sets of standardized single word reading and literacy tests:

1. Woodcock-Johnson IV (WJ-IV) subtests: untimed word identification (word reading accuracy) and word attack (nonword reading accuracy), as well as tests for three different aspects of phonological awareness: sound blending, segmentation, and sound awareness.^65^ We found that as a result of school-specific intervention protocols, our DD cohort performed above 50% of age-matched peers on two of these tests (see Table 1).
2. The Timed Test of One-Word Reading Efficiency, version 2 (TOWRE-2), which has two subtests, one measuring sight word recognition efficiency based on timed sight-word reading efficiency (SWE), and one measuring phonemic decoding efficiency (PDE) based on timed non-word reading efficiency.^66^

Of note, most DD children also completed the Gray Oral Reading Test, version 5 (GORT), which assesses oral reading fluency and comprehension based on passages and stories reading.^67^ Because this test assessed complex reading comprehension rather than phonological decoding or single word reading, we do not include it in any of our main analyses. However, scores are reported in supplements for completeness (Supplementary Table 2).

Due to limitations of time, protocol updates, or subject fatigue, not all DD children were able to complete all of the tasks (see Table 1 for sample size details of each test). TD children participated in an abbreviated study protocol and completed only matrix reasoning, digit span forward and backward, rapid picture naming, and TOWRE-2 tests.

### Auditory processing tasks

#### Non-speech amplitude rise time task (ART)

We evaluated the perceptual threshold for amplitude rise time with a standard adaptive staircase procedure, using a 3-steps-down 1-step-up procedure converging to a 79% just noticeable difference (JND).^68^ The perceptual acuity was measured by using a two-alternative forced-choice (2AFC) design, namely in each trial, participants heard two harmonic tones with a triangular amplitude shape and were asked to identify which of the two tones had a longer rise time (softer onset). Tone rise times on subsequent trials were adjusted according to a child’s response: it was increased following an incorrect response and decreased after a series of three consecutive correct responses. The standard tone rise time was fixed to 15 ms, whereas the test tone had an initial rise time of 300 ms, varying between 15 and 500 ms. The inter-stimulus interval was fixed to 350 ms. The task terminated after eight response reversals (i.e., switches between correct and incorrect responses) or the maximum possible 80 trials.^9^ To account for worse overall performance in children, we defined successful completion of the task as performance above an accuracy criterion of 65 % (see Supplementary Table 1).^69,70^ The rise time JND was then calculated as the average rise time on the last 8 reversal trials. A lower raw rise time JND indicates better amplitude rise time performance. To best evaluate amplitude rise time abilities in the 2AFC design, we also calculated the accuracy of the reference stimulus in the first and the second interval, referring to Raviv *et al.*^71^. In line with previous work, the accuracy of the reference in the first interval was higher than that of the second interval across groups, while no significant group differences were found in the pattern of differences between the first and second reference intervals. For further analyses, raw JNDs were z-scored and inverted such that higher z-scores indicate better performance.

#### Speech in noise task (SiN)

Speech in noise perception accuracy was tested using the single syllable in background noise. On each trial, children heard a single syllable and were asked to repeat what they heard. The examiner recorded the responses. Syllables were consonant-vowel combinations, namely 12 consonants covering three phonetic features (voicing, place, and manner) in English and ending with the vowel /a/. Each syllable was repeated 5 times and presented in two noise conditions, at −6 and −12dB relative to the noise level. Noise conditions were blocked, with the 6 dB condition administered first. Before the noise conditions, all syllables were presented once in quiet, in a practice block. We calculated the percentage of correct responses for each syllable at each noise level. In addition, we examined confusion patterns on error trials to evaluate the percentage of transmitted information for the place, manner, and voicing of articulation.^20,72^ Notably, all raw scores were converted to standardized z-scores for further analyses.

### Image acquisition and processing

Neuroimaging data were acquired with a 3.0 Tesla Siemens Prisma MRI scanner. T1-weighted (T1w) three-dimensional sagittal Magnetization Prepared Rapid Acquisition Gradient Echo (MPRAGE) images were acquired with the following parameters: TR = 2300 ms, TE = 2.98 ms, TI = 900 ms, flip angle = 9°, field of view (FOV) = 256 × 240 × 160 mm^3^, spatial resolution = 1 × 1 × 1 mm^3^, parallel imaging acceleration factor (iPAT) = 2.

T1w images were preprocessed using the FreeSurfer toolbox (version 6.0.0) for cortical reconstruction and volumetric segmentation. Once surfaces were reconstructed, an array of anatomical measures, including cortical thickness, surface area, and local gyrification index (LGI), were then automatically calculated at each vertex of the cortex. The LGI, a unitless measure quantifying gyrification of the brain was investigated in the current study, as a metric of the amount of cortex buried within the sulcal folds as compared with the amount of cortex on the outer visible cortex. A large gyrification index indicates a cortex with extensive folding and a small gyrification index indicates a cortex with limited folding. The vertex-wise maps of individuals were aligned to the FreeSurfer *fsaverage* surface-based template and smoothed using a 5 mm FWHM Gaussian kernel for group analysis (see detailed image processing in the Supplementary Methods).^73^

### Statistical analysis

#### Behavioral analyses of auditory and language tasks

First, we tested for group-level differences between DD and TD groups’ performance on the ART and SiN tasks using Welch’s two-sample *t-test*, which accounts for unequal sample sizes between groups.

Pairwise associations between reading measures and auditory tasks were evaluated using Pearson’s correlation. We further evaluated the predictive relationship of auditory processing abilities on reading and phonology through hierarchical regression analyses. Models involved reading, phonological awareness, and other phonological measures, including digit spans (as a proxy of phonological memory), and rapid picture naming (as a proxy of rapid automatized naming) as dependent variables. Covariates such as age and sex, matrix reasoning, and ROWPVT vocabulary were included in the model, with auditory processing scores as the independent variables. The full model was: reading ∼ age + sex + matrix reasoning + ROWPVT vocabulary (Step 1) + auditory ability (Step 2). This analysis was performed separately for each of the auditory processing tasks. Due to the small sample size and subset of behavioral data available in the TD group, we performed the above analyses within the DD group. If not specified, all analyses within the DD controlled for age and sex, with matrix reasoning additionally controlled for in the group comparisons between DD and TD to rule out that the observed effects were driven by trend-level differences in this task. All *p-values* are two-tailed with a threshold of *p* < 0.05. Additionally, all behavioral scores are percentile scores and auditory processing scores are z-transformed values.

In addition to the regression analysis, we analyzed the behavioral data using direct acyclic graphs (DAGs) via the *dagitty package* in R to conceptualize the causal impacts of auditory abilities on reading through phonological measures between multiple sets of behavioral variables.^74,75^ Auditory measures were considered as exposures, with reading measures as outcomes, and phonological measures, including phonological awareness (sound awareness, sound blending, and segmentation), rapid picture naming, and digit spans as mediators. Age, sex, ROWPVT vocabulary, and matrix reasoning were accounted for as confounders (see Supplementary Fig. 1 for the conceptual models and Supplementary Results for details). These DAGs informed the structural equation modeling (SEM) via the *lavaan package* in R to statistically evaluate the hypothesized causal paths and the mediation effects.^76,77^ To address sample size constraints and ensure robust estimation, we simplified the model by considering only phonological awareness as the mediator in the main text, using residuals obtained after regressing out confounders as inputs (see Supplementary Results for other phonological measures).

#### Brain-behavioral correlations

We tested whether performance on the auditory processing tasks was correlated with local cortical gyrification. This analysis was performed at the whole-brain level for the DD group and for the whole cohort. Age, sex, and total brain volume were included as covariates of no interest in all analyses. A common threshold for surface-based analysis was used with a cluster-forming threshold of *p-value* < 0.005 at a cluster level of *p-value* < 0.05 corrected for multiple comparisons based on random field theory.^78–80^ All brain-behavior correlations were performed using the s*urfstat* toolbox implemented in MATLAB.

### Data availability

The data that support the findings of this study are available on request from the corresponding and senior authors. The data are not publicly available due to limitations of our ethics approval. Data requests can be submitted at: https://memory.ucsf.edu/research-trials/professional/open-science. Following a UCSF-regulated procedure, access will be granted to designated individuals in line with ethical guidelines on the reuse of sensitive data. This would require the submission of a Material Transfer Agreement. Commercial use will not be approved.

## Results

### Left superior temporal gyrus underlies different auditory processing in children

One hundred and ten participants, including 78 children with DD and 32 TD children, who successfully completed at least one of the auditory tasks were included in the present study (for detailed demographics and behavioral characteristics see Table 1). We first compared sensitivity to non-speech amplitude modulations, evaluated by a non-speech amplitude rise time (ART) task and a speech in noise (SiN) task, between DD and TD groups.

Amplitude rise time discrimination was impaired in DD (*mean*_rawJND_ = 226.57, *SD* = 155.51), evident in significantly elevated thresholds in this group as compared to TD (*mean*_rawJND_ = 127.81, *SD* = 84.28; *welch’s-t*(70.74) = −3.03, *p* < 0.01, Figure 1A). In contrast, groups did not differ in the SiN task (main effect of group: *F*(1,167) = 1.97, *p* = 0.16; group by noise interaction effect: *F*(1, 167) = 0.91, *p* = 0.34), with overall more impaired performance at higher relative noise levels (group average: 6dB: *mean* = 43.50%, 12 dB: *mean* = 25.00%, see Figure 1B) in both groups (main effect of noise level: *F*(1,167) = 253.10, *p* < 0.01). Given the very low accuracy in the 12 dB condition, we focused on the 6 dB condition in all subsequent analyses (results for the 12 dB condition see Supplementary Fig. 2,3, and the Supplementary Results). As previous work showed selective impairments in DD for the perception of certain consonant types, we also analyzed recognition accuracy for single phonetic features.^20^ Noise differentially affected phonetic features in both groups (main effect of feature: *F*(1,268) = 8.92, *p* < 0.01), but without differences between groups (main effect of group: *F*(1,268) = 0.05, *p* = 0.83; group by phonetic features interaction effect: *F*(1,268) = 0.24, *p* = 0.78). Importantly, behavioral performance was not correlated between the two tasks either in DD (*r* = −0.07, *p* = 0.63, Figure 2A) or across groups. This supports the dissociation between amplitude modulation and speech perception abilities and suggests that each might contribute to different aspects of speech processing during development.

**Figure 1.**
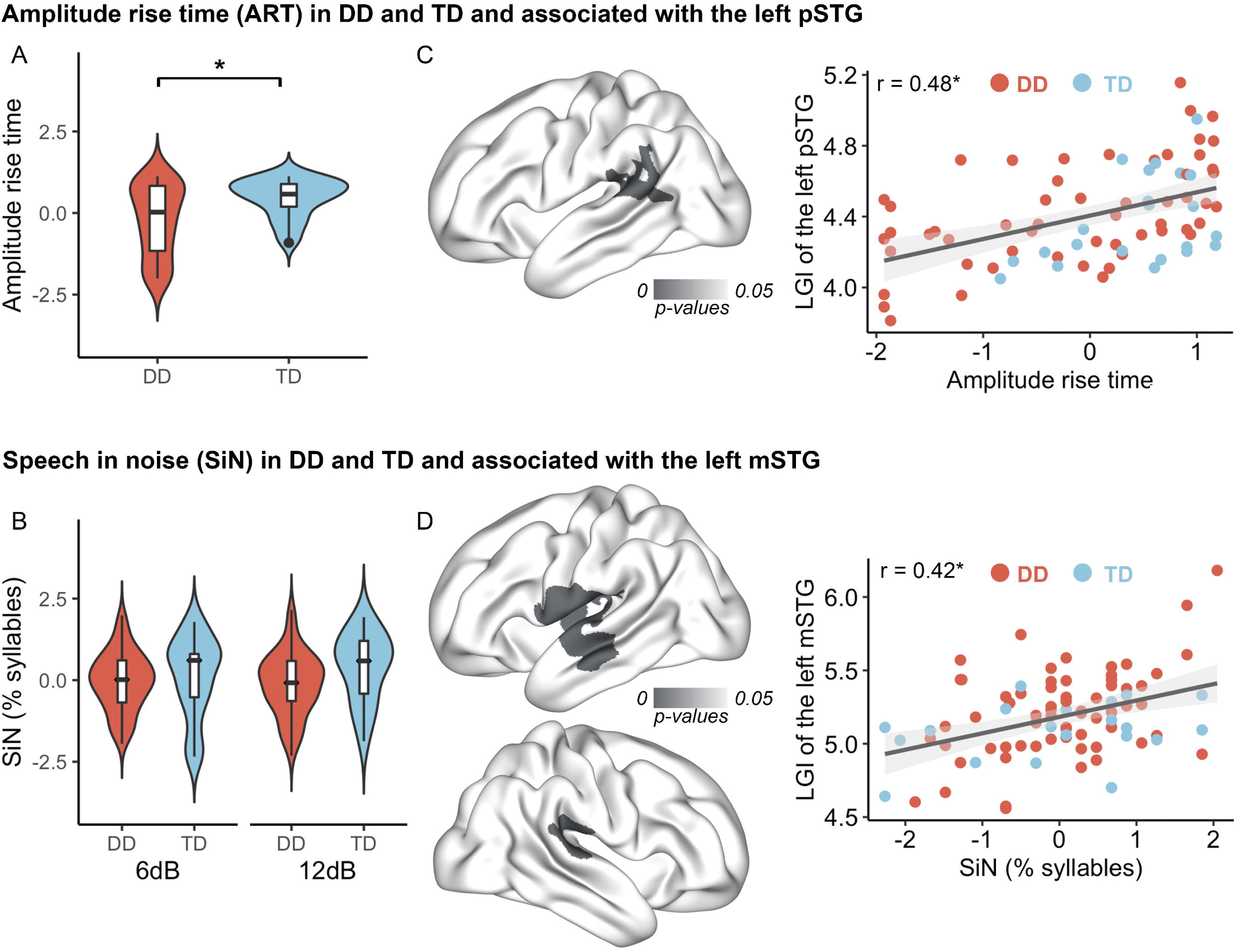
Auditory processing abilities in children with DD compared to TD and their relationship with cortical folding in the whole brain. (**A**) Children with DD showed decreased amplitude rise time discrimination abilities. (**B**) No significant group difference in speech in noise recognition abilities between DD and TD groups at either level of noise. (**C**) Amplitude rise time discrimination abilities were associated with the local gyrification index (LGI) of the left posterior superior temporal gyrus (pSTG) (*p* < 0.05 and FWE-corrected). (**D**) Speech in noise recognition abilities were correlated with the LGI in the left middle STG (mSTG) (*p* < 0.05 and FWE-corrected). Red and blue denote DD and TD groups, separately. All auditory processing ability raw scores were converted to standardized z-scores (see Table 1 for raw scores). * *P* < 0.05.

**Figure 2.**
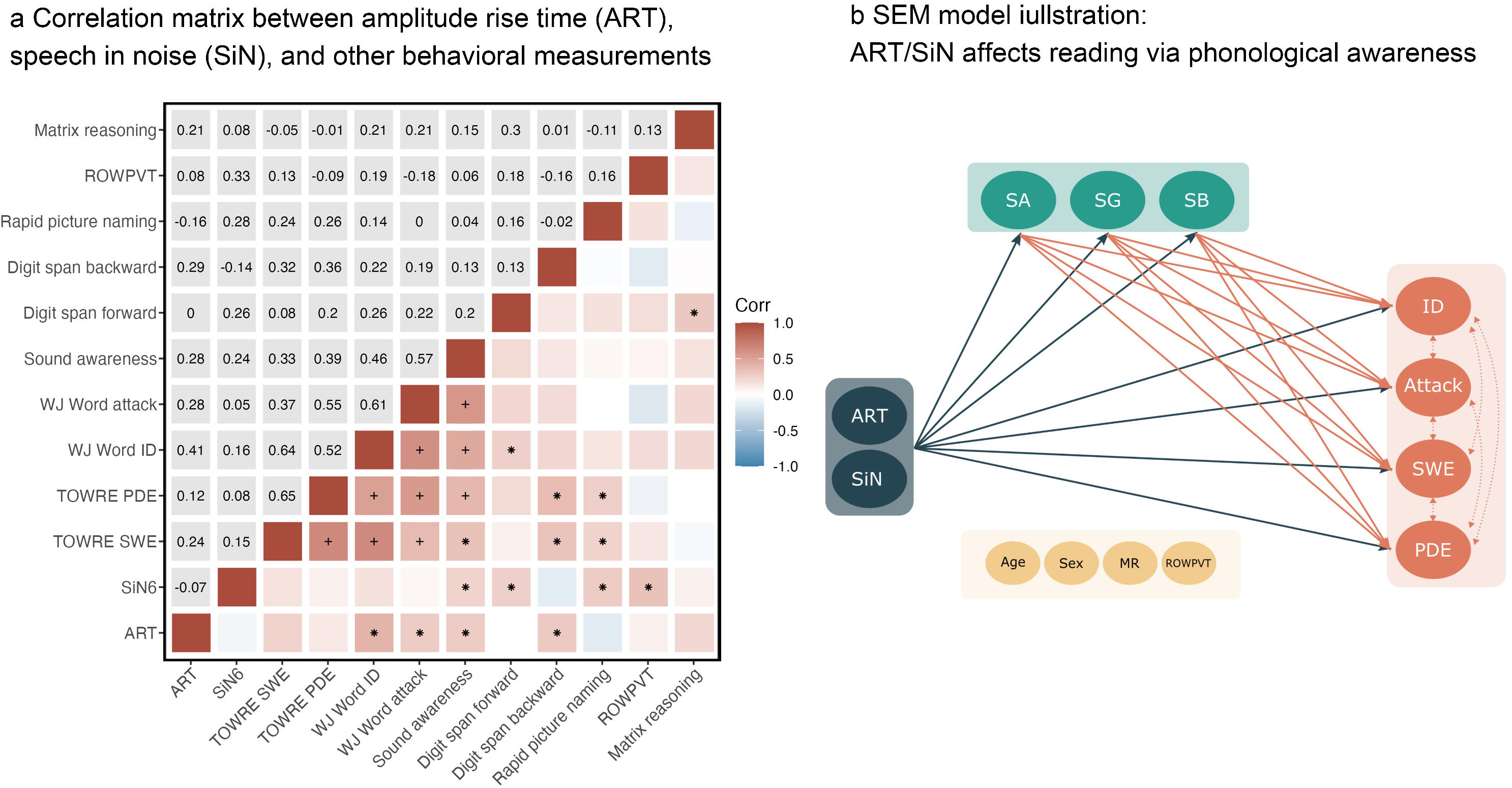
Amplitude rise time is predictive of reading and speech in noise is predictive of phonological sound awareness in children with DD. (**A**) Correlation matrix between amplitude rise time, speech in noise, reading, phonology, and cognitive measures. The upper panel illustrates the correlation coefficients after controlling for age and sex, and the lower panel illustrates the pattern of correlations. (**B**) Schematic of structural equation modeling (SEM) illustrating the mediation effects of phonological awareness on the influence of auditory processing abilities on reading. ART: amplitude rise time, SiN: speech in noise, ROWPVT: receptive one-word picture vocabulary test, MR: matrix reasoning, ID: WJ word identification, Attack: WJ word attack, SWE: TOWRE single-word reading efficiency, PDE: TOWRE phonemic decoding efficiency, SA: sound awareness, SG: segmentation, SB: sound blending. + *P* < 0.05 after multiple comparisons correction, * *P* < 0.05,. *P* < 0.10.

Next, we investigated how variation in cortical structure in posterior and middle STG relates to perception of amplitude modulations and to perception of speech in noise. Whole-brain LGI and behavior correlation analyses showed that better amplitude rise time discrimination was associated with greater cortical folding in the left pSTG (*r* = 0.48, *p* < 0.01, FWE corrected, Figure 1C) across both groups (*n* = 78, DD/TD = 56/22). Likewise, speech in noise task performance was correlated with LGI in the left mid-anterior STG (*r* = 0.42, *p* < 0.01, *n* = 84, DD/TD = 60/24). Additionally, speech in noise task performance was also correlated with LGI in clusters in the left insula, precentral gyrus (*p* < 0.01), and the right pSTG (*p* < 0.01, all FWE corrected, Figure 1D), suggesting that these areas might be part of the network involved in speech comprehension under challenging listening conditions. Of note, similar results were also observed within the DD group for each of the auditory tasks. Group comparisons between LGI in the DD and TD groups showed no differences within the identified clusters, nor at the whole-brain level.

Crucially, we found a high overlap between the two significant left STG clusters and functional zones previously identified by iEEG in the left STG (Figure 3).^11,55,56^ The significant rise time cluster overlapped with the speech onset zone in pSTG, while the speech in noise cluster overlapped with the phonetic features zone in mSTG. Overall, these analyses showed that cortical folding of different functional subdivisions of the STG was related to distinct aspects of auditory processing in children with and without DD.

**Figure 3.**
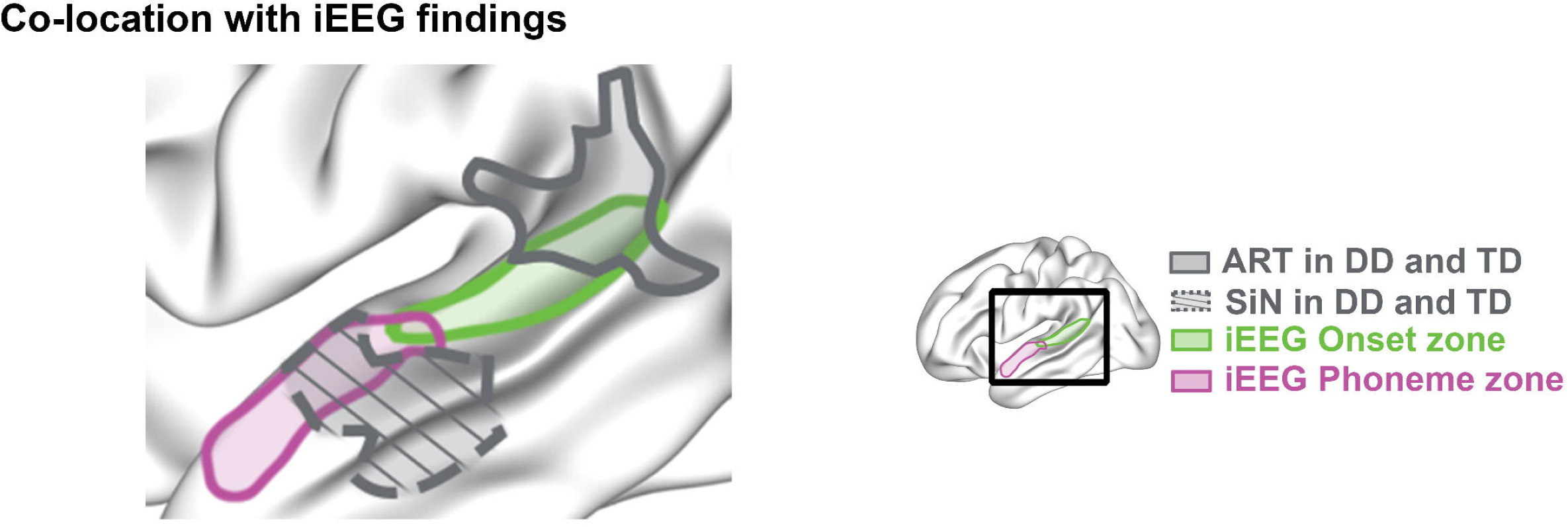
The left pSTG and mSTG fall in the speech onset and phonetic feature zone defined in our iEEG work, respectively. Figure adapted from Oganian *et al*.^11^. Notes: speech onset is color-coded in green and phonetic feature zone is color-coded in purple. The solid and dashed gray blobs denote the left pSTG associated with amplitude rise time and the left mSTG associated with speech in noise perception.

### Amplitude rise time is associated with reading and phonological abilities in DD

Next, we aimed to understand how the two auditory tasks are related to the main deficits in reading and phonology in DD. Based on prior literature and the brain-behavior correlations in our data, we hypothesized that amplitude rise time performance would be related to reading performance, whereas speech in noise performance might be more relevant for phonological awareness.^11,55,56,81^ Children with DD underwent a comprehensive battery of neuropsychological and academic testing. Due to the small sample size and subset of behavioral data available in the TD group, we conducted all behavioral analyses within the DD group. Figure 2A illustrates the correlation pattern across variables. Given the well-established, complex pattern of dependencies between those measures, we used hierarchical regression analysis to investigate how amplitude rise time discrimination predicts multiple reading and phonological measures (phonological awareness, rapid picture naming, and digit spans), accounting for age, sex, matrix reasoning, and ROWPVT vocabulary. Results indicated that rise time predicts WJ word ID (ΔR^2^ = 0.11, *t* = 2.74, uncorrected *p* < 0.01, β *=* 0.35), sound blending (ΔR^2^ = 0.07, *t* = 2.43, uncorrected *p* = 0.02, β *=* 0.30), and sound segmentation (ΔR^2^ = 0.13, *t* = 3.39, uncorrected *p* < 0.01, β *=* 0.38) (see Supplementary Table 3 for all hierarchical regressions of reading and phonological scores in DD).

We then used structural equation modeling (SEM) to examine whether amplitude rise time discrimination influences reading measures through phonological awareness, controlling for confounders. The SEM results indicated a good model fit (*chi-square p* = 0.46, see Table 2, Figure 2B, and Supplementary Table 4), with significant direct effects of rise time on segmentation, sound blending, and real word reading measures (WJ word ID: *p* = 0.05, TOWRE SWE: *p* < 0.05). Phonological awareness significantly influenced reading, with sound awareness affecting all reading measures, segmentation specifically influencing TOWRE SWE, and sound blending specifically influencing WJ word attack (*ps.* < 0.05, Supplementary Table 4). Notably, segmentation mediated the effects of rise time on TOWRE SWE (*p* < 0.05) and sound blending marginally mediated the effects of rise time on WJ word attack (*p* = 0.08). Total effects involving amplitude rise time were particularly noted on real word reading measures (WJ word ID: *p* < 0.01, TOWRE SWE: *p* = 0.09, *trend-level*). These results show that amplitude rise time directly impacts reading and is mediated by segmentation abilities. Taken together, our behavioral analyses suggested that children with DD are impaired in their perception of amplitude rise time, independent of general cognitive ability (β = 0.24, *t* = 1.49, *p* = 0.14), vocabulary (β = 0.05, *t* = 0.35, *p* = 0.73), as well as sex (β = −0.13, *t* = −0.44, *p* = 0.66) and age (β = 0.22, *t* = 1.45, *p* = 0.16). Importantly, these results confirm that amplitude rise time discrimination is a major contributing factor to DD’s impaired reading abilities, notably real word reading.

**Table 2.**
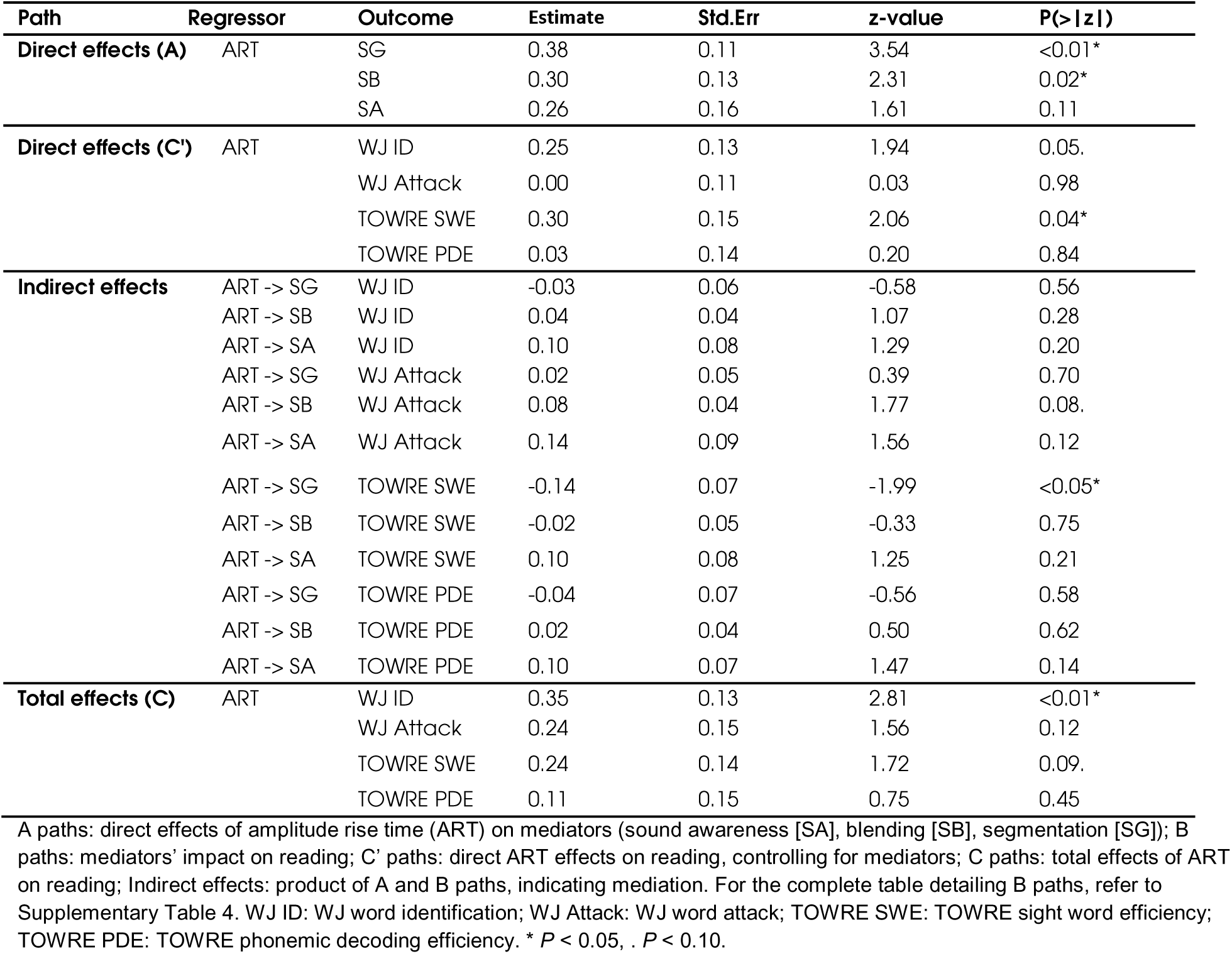
SEM Overview: amplitude rise time (ART) contributes to reading via phonological awareness.

### Speech in noise is associated with phonological abilities in DD

Behavior in the speech in noise task was correlated with cortical gyrification in middle STG, the main cortical region representing phonetic and phonological speech content.^11,55,56^ Thus, we hypothesized that performance on this task would predict phonological awareness in the DD group. In the hierarchical regression model, including age, sex, matrix reasoning, ROWPVT vocabulary, speech in noise perception accounted only for sound awareness (ΔR^2^ = 0.05, *t* = 2.09, *p* < 0.05, β *=* 0.29), but not other phonological or reading measures (see Supplementary Table 5). Subsequent SEM analysis indicated a poor model fit (*chi-square p* < 0.01), suggesting caution in interpreting the path coefficients. Specifically, speech in noise showed a significant direct effect on sound awareness (*p* = 0.01), but not on other phonological or reading measures. Indirect effects on several reading measures through sound awareness were demonstrated (see Supplementary Table 6). However, neither total nor direct effects were significant, highlighting a less direct influence on reading. Collectively, SEM results support that speech in noise recognition, influenced by vocabulary (β = 0.32, *t* = 2.71, *p* < 0.01) and age (β = 0.42, *t* = 3.22, *p* < 0.01) but independent of general cognitive ability (β = 0.06, *t* = 0.41, *p* = 0.68) and sex (β = −0.31, *t* = −1.20, *p* = 0.23), is associated with phonological sound awareness in children with DD.

## Discussion

We provide a neuroanatomical and behavioral dissociation between non-speech auditory processing of sound amplitude rises and speech recognition abilities in children with developmental dyslexia (DD) and a neurotypical control group. Behaviorally, DD were impaired in non-speech amplitude rise time discrimination, but not in speech in noise recognition. Neurally, cortical LGI differentiates the neural substrates related to the two tasks. Namely, across groups, amplitude rise time discrimination was positively correlated with LGI of left pSTG, whereas speech in noise perception was positively correlated with LGI of left mSTG. This dissociation was further manifested in distinct association patterns with reading and phonology between the two tasks in DD.

The observed dissociation between non-speech amplitude rise time and speech in noise perception is well aligned with recently discovered distinct response profiles in posterior and middle STG in iEEG recordings, enabled by a close match between the stimuli used in our amplitude rise time task and those used to study neural processing of amplitude rises with iEEG (see Figure 3).^11,55,56^ The observed correlation between gyrification and behavioral performance in our cohort is in line with prior studies that argue in favor of a functional gradient for processing speech sounds along the posterior-to-anterior axis of the STG.^31^ We extend these results to the perspective of DD: psychophysics results along with the replicated spatial distributions across studies emphasize the idea that anterior and posterior STG are specialized for different processes during speech perception. Although gyrification did not differ between the dyslexic cohort and controls, our study aligns with prior work in DD using MRI that identified atypical STG activation in a range of auditory and phonetic tasks.^30–32,59^ Specifically, our findings are in line with earlier neurophysiological studies that reported atypical speech in noise processing and cortical tracking of speech in the bilateral mid-superior temporal gyrus in dyslexic children.^82^ Furthermore, other MEG studies also identified amplitude envelope processing impairments in the STG during naturalistic story-listening in children with DD.^18^ Interestingly, enhancing speech edges has also been shown to significantly improve neural processing in children with dyslexia, as evidenced by stronger speech tracking in the delta band within the STG.^83^ Additionally, atypical cortical morphometry, e.g., thickness and surface area, has been reported in temporal cortex.^36,38,52^ Our findings also add to previous literature reporting atypical cortical folding in the occipitotemporal and temporoparietal cortices in DD.^43,45,84^ More importantly, our results suggest that distinct aspects of the STG support phonological and non-speech auditory processing and highlight the distinct roles of posterior and middle STG in speech sound processing in developmental populations.

In our cohort brain to behavior correlations were consistent across groups with no LGI differences between DD and TD children. This stands in contrast to previously reported structural alteration in the temporal cortex in DD.^34,36,38,45^ Particularly, prior studies used a range of different folding-related indices reflecting different geometric properties of the cortex (e.g., folding index and mean curvature),^85–87^ with only some reporting folding-related alterations in DD.^43–45,84^ The absence of group-level differences in our study may be due to the small size of our TD group alongside the large variation in the DD group itself. The overall similar correlational pattern in TD and DD might also indicate that underlying deficits in DD lead to overall reduced LGI but do not alter the role of the STG in sound processing. Further studies including longitudinal data will be necessary to further clarify the relationship between auditory processing, phoneme representations and the development of cortical folding of the STG.

As expected, we found that children with DD were impaired in amplitude rise time discrimination. In fact, rise time discrimination deficits in children and adults with DD are well-documented and among the most robust auditory deficits in DD.^8,15,17^ Prior studies also found that rise time deficits were related to reading and phonological awareness.^14,17,88,89^ Our results are in line with those studies, showing that rise time deficits are direct predictors of reading abilities, particularly real word reading. Intriguingly, amplitude rise time discrimination deficits, present from newborn age in infants at familial risk for dyslexia, were not correlated with age, supporting a precursor role of amplitude rise time for phonological and reading skills in DD, as previously proposed.^12,21,22,90–92^ Finally, we want to note that recent studies showed genetic correlations between neuroanatomy of the left posterior superior temporal cortex, where onset rise time is processed, and reading and language measures in cohorts including young children as well as young adults, which might further point to the heritable nature of the role of auditory processing abilities in speech and reading skills.^93^

Speech in noise perception was not impaired in our cohort of children with DD but was predictive of phonological sound awareness.^94^ Together with the impairment in amplitude rise time discrimination, this aligns with previous studies that showed dissociations between different auditory perception tasks in DD.^2^ Beyond that, prior findings on speech in noise perception in DD have been mixed [impaired: ^12,20,95,96^; intact, particularly in adults: ^81,97,98^]. Interestingly, changes in speech in noise among individuals with DD were not observed in a study comparing them to controls matched for reading level.^82^ This implies that these alterations may simply be influenced by diminished reading exposure rather than serving as a root cause of dyslexia. In fact, it has been previously suggested that speech in noise abilities improve substantially with age and with reading instruction and phonological awareness and might not persist into adulthood.^12,97,99^ Together with the correlation between phonological sound awareness and speech in noise perception, it is possible that our participants overcompensated their speech in noise deficits. This may particularly be the case as our cohort of DD received targeted interventions with phonological awareness training. This account is supported by the better speech in noise perception in older participants - those who had more reading and phonological training. It is further supported by neuroimaging studies that found a compensatory role of right STG in speech in noise perception in adults with DD.^97^ This is in line with higher LGI of the right STG with better speech in noise recognition in our cohort. In addition to age-related increases, our findings showed that vocabulary accounted for speech in noise performance, suggesting that speech in noise deficits reflect a complex profile beyond dyslexia.^100,101^ Differences in subgroups, based on speech in noise tasks, verified this complex relationship (see Supplementary Table 7). Overall, our results suggest that speech perception deficits are related to phonological processing skills in DD but, unlike deficits with amplitude rise time perception, are not directly related to reading abilities.

Amplitude rise time discrimination is distinct from speech in noise perception in behavior and in neuroanatomy. This raises the possibility that performance on these tasks might define distinct phenotypes in DD, each characterized by distinct patterns of severity in auditory deficits (see Supplementary Table 7 and Supplementary Results). However, as stated above, amplitude rise time deficits are present in newborns at familial risk for dyslexia, whereas deficits in speech in noise are not observed until preschool age, and continue to improve dramatically with age until late childhood.^12,21^ This again favors the hypothesis of potential linguistic compensation for speech in noise deficits during development.^81,99,102^ Taken together with the different neural correlations between the tasks, our results emphasize that the specific expression of auditory deficits differs between individuals, with different auditory impairments affecting phoneme awareness and reading. The distinct neural correlates of our two specific tasks highlight that these differences also come hand in hand with distinct neural impairments. Regardless of the existence of distinct “auditory” DD phenotypes, a neuroanatomical dissociation between different auditory processing abilities is demonstrated here, which is critical for clinical applications, and might provide a framework for the design and evaluation of differential approaches to interventions for DD. Future longitudinal studies should follow younger children with a broader range of tasks to investigate associations between different auditory abilities and their relation to neural development along the STG in neurotypical and DD populations. Although the observed amplitude rise time deficits support the auditory processing deficits theory in DD, it does not refute other additional sources of deficits in DD, which were not investigated in the current study.^2^ Indeed, given the large range of performance on the auditory tasks within the DD group, it is possible that additional cognitive and neuroanatomical mechanisms also contributed to the reading challenges in this cohort. Indeed, the heterogeneity of deficits in DD, which may not be limited to auditory processing deficits, further stresses the importance of studying individual and group differences in clinical populations.

Several limitations should be considered when interpreting our results. First, our study included a relatively small cohort of TD children. As discussed above, null results at the group level in behavior and neural analyses warrant future studies with a larger TD group. However, considering the challenges of collecting large and multimodal data in clinical neurodevelopmental populations, our cohort of 110 children is still above the average sample size in the field. Next, although LGI is considered to be one of the most sensitive neural measures to distinguish DD and TD, it is largely under-investigated compared to other cortical geometric properties.^43^ In particular, little is known about whether LGI relates to other neurodevelopmental changes, such as the commonly investigated cortical thickness, surface area, and myelination.^73,85^ This highlights the need for investigations of the underlying biological mechanism from different perspectives. Our data showed that auditory processing abilities may show differential sensitivity to age. While we have included a wide age range of school-age children, future studies should consider incorporating younger children to better understand auditory processing and its neural mechanism developmentally. Beyond that, the correlational nature of the current study constrains us from establishing causality between the abnormalities of the brain and behavior, emphasizing the need for future research.

Overall, we provide the first evidence for distinct contributions of posterior and middle STG to different auditory processing deficits in DD. Our study enhances the understanding of auditory processing deficits in DD by characterizing how distinct auditory tasks are related to reading, phonology, and cortical neuroanatomy. Our results show that auditory and phonological processing difficulties may arise through multiple underlying mechanisms, which vary across individuals. Possible clinical implications of this pattern call for future studies on the inter-individual variability in DD phenotypes and their response to interventions.

## Acknowledgements

We would like to express our sincere thanks to the parents and children who participated in this project. We are also grateful to the anonymous reviewers for their careful reading of our manuscript and their constructive comments.

## Supplementary material

Supplementary material is available at Brain online.

