## Supplementary materials for "Anatomical and behavioral correlates of auditory perception in developmental dyslexia"

### Supplementary material

#### Supplementary Methods

##### Participants

Initially, one hundred and thirty-five participants, including 90 children with developmental dyslexia (DD) and 45 typically developing (TD) children, who completed the auditory tasks, were included in the present study. Inclusion criteria based on the behavioral data were as follows: 1) Matrix reasoning falls in and above the normal range ($16^{th}$ percentile); 2) to account for worse overall performance in children on the amplitude rise time (ART) task, only participants with accuracy above 65% were included in the analysis; 3) for the speech in noise task (SiN), only children with performance within the 1.5 interquartile range (IQR) were included in the analysis. See Supplementary Table 1 for more details.

##### Image processing

Surfaced-based preprocessing: Before preprocessing, T1w images were visually inspected for potential artifacts caused by head motion to ensure that brain tissues could be well-differentiated. Cortical reconstruction and volumetric segmentation were performed using the FreeSurfer toolbox (version 6.0.0). The reconstructed surfaces were visually inspected and manually edited for inaccuracies. The vertex-wise maps of individuals were aligned to the FreeSurfer *fsaverage* surface-based template, which has been shown that using for children from ages 4 to 11 years does not result in an age-associated bias.[1]

### Supplementary Results

#### Neuropsychological and academic assessments

##### Other (passages and stories reading) diagnostic reading measures

Supplementary Table 2 reports reading scores for the Gray Oral Reading Test, version 5 [GORT[2]], which assesses oral reading fluency and comprehension based on passages and stories reading. The fact that the tests were administered to only very few TD children, along with the unequal sample sizes between the two groups, largely constrained the statistical power of group comparisons, which showed only a weak trend towards group differences.

##### Principal component analysis (PCA) for the four reading scores

Instead of correlating auditory processing abilities with multiple reading scores, we used data-driven PCA (R package "*factoextra*") to reduce the four single-word reading scores to two unbiased principal components (PCs), together accounting for 85.26% (70.52% for PC1 and 14.74% for PC2) of the total variance. Specifically, WJ word ID, TOWRE SWE, WJ word attack, and TOWRE PDE account for 27.79, 24.15, 22.23, and 25.83% of the total variance of PC1, and account for 1.43, 43.47, 54.51, and 0.59% of the PC2. Supplementary Fig. 2 presents pairwise correlations between auditory processing abilities, reading (including two reading PCs), phonological, and other cognitive abilities for DD children. The correlation patterns based on the PCs qualitatively follow the results of correlations for single reading scores and amplitude rise time abilities (PC1: $r = 0.32$, $p = 0.02$; PC2: $r = 0.08$, $p = 0.54$) in DD. Speech in noise recognition abilities were not correlated with any of the reading measures and reading PCs.

### Speech in noise (SiN) task

#### Marginally impaired SiN recognition abilities at 12 dB in DD compared to TD group

In the main text, we showed that comprehension was impaired at higher relative noise levels in both groups (main effect of noise level: $F(1,167) = 253.10$, $p < 0.01$). Comparing the performance in the 12 dB condition between DD and TD groups, we observed a marginal group difference in the accuracy of SiN recognition ($p = 0.08$, $F(1,77) = 3.15$, *mean* = 23.07% and 26.94% for DD and TD, respectively, Supplementary Fig. 3A).

We continued to examine the recognition accuracy for single phonetic features as previous work showed selective impairments in DD for the perception of certain consonant types.[3] We observed significant differences between groups and between features (main effect of feature: $p < 0.01$, $F(1,235) = 42.28$; main effect of group: $p = 0.02$, $F(1,235) = 5.94$; interaction effect: $p = 0.94$, $F(1,235) = 0.06$). This reflects that the overall performance on phonetic feature recognition in children with DD (*mean* = 56.42%) was impaired compared to TD (59.69%). Across two groups, the recognition of place of articulation (52.40%) was much more difficult than the recognition of voicing (57.40%) and manner of articulation (64.30%). Post-hoc analyses further confirmed a marginal difference in place-of-articulation perception between groups ($p = 0.06$, $F(1,77) = 3.58$; manner of articulation: $p = 0.18$, $F(1,77) = 1.87$; voicing: $p = 0.32$, $F(1,77) = 1.02$), indicating a trend-level selective impairment for the perception of place of articulation of consonants in DD (Supplementary Fig. 3B).

Pairwise correlations (Supplementary Fig. 2) show that speech in noise recognition abilities at 12 dB were positively correlated with digit span forward ($r = 0.30$, $p = 0.02$) and WJ word attack in the DD group ($r = 0.25$, $p = 0.05$), but not with phonological sound awareness, matrix reasoning, and age ($r = 0.05$, $p = 0.71$). Speech in noise recognition abilities at 12 dB were also not correlated with the amplitude rise time ($r = -0.20$, $p = 0.19$) and the speech in noise performance at 6 dB ($r = 0.11$, $p = 0.42$).

**Using direct acyclic graphs (DAGs) to conceptualize the causal impacts of auditory abilities on reading through phonological processing abilities**

Building on regression analysis, we constructed direct acyclic graphs (DAGs) using *dagitty package* in R to conceptualize the causal impacts of auditory abilities on reading through phonological awareness.[4,5] A broader range of variables were considered in the DAGs. Auditory measures were defined as exposure, multiple reading measures as outcomes, and phonological measures, including phonological awareness (sound awareness: SA, sound blending: SB, segmentation: SG), rapid picture naming (a proxy of rapid automatized naming, RAN), and digit spans (as a proxy of phonological memory) as mediators. Age, sex, receptive one-word picture vocabulary (ROWPVT) and matrix reasoning (MR) were considered as confounders (see Supplementary Fig. 1). We evaluated the consistency between the conceptual DAGs and the dataset applying the d-separation criterion to check for implied conditional independencies. Each independence was tested for zero correlation, where a *p-value* greater than 0.05 typically indicates that the conditional independence holds true in the data.[4,5] It showed that this initial DAG did not align with the data due to the moderate correlation among the four reading measures. We refined the DAG by adding direct paths between reading outcomes to better reflect the theoretical interrelations and to satisfy consistency between the adjusted DAG and the dataset. This refined model was then tested using structural equation modeling (SEM) to provide the statistical inferences of the causal pathways influencing reading abilities.

In addition to exploring the mediating effect of phonological awareness in the main text, we incorporated both rise time and speech in noise into one model to more comprehensively examine how auditory processing abilities influence reading. The SEM results indicated a good model fit (*chi-square* $p = 0.36$, see Supplementary Table 8 for details). Specifically, amplitude rise time showed significant direct effects on segmentation, sound blending, TOWRE SWE (*ps.* $< 0.05$), and marginally on WJ word ID ($p = 0.05$). However, the mediation effects of auditory processing abilities on reading were generally not significant, except for a notable effect of rise time on TOWRE SWE through segmentation ($p < 0.05$). Total effects were noted only for the impact of amplitude rise time on real word reading measures, i.e., WJ word ID ($p = 0.01$). Speech in noise showed a significant direct effect on sound awareness ($p = 0.01$), which in turn mediated the relationship of speech in noise on WJ word ID ($p = 0.03$) and on WJ work attack ($p = 0.04$). However, neither total nor direct effects were significant, highlighting a less direct influence on reading. Overall, this complementary analysis delineates the different roles of amplitude rise time and speech in noise in reading outcomes, with rise time more directly influencing real word reading and speech in noise more directly affecting phonological sound awareness.

Furthermore, add to the mediating effect of phonological awareness in the main text, we tested whether other phonological processing abilities, specifically phonological memory and RAN, also mediated the relationship between auditory processing abilities and reading. We performed separate SEM analyses with phonological memory and RAN as mediators. For the relationship between amplitude rise time to reading, the SEM model with phonological memory, namely digit span forward (DSF) and digit span backward (DSB) as mediators, indicated a good fit (*chi-square* $p = 0.56$, see Supplementary Table 9). Amplitude rise time exhibited a significant direct effect on DSB ($p = 0.04$), and DSB directly influenced TOWRE SWE ($p = 0.08$) and TOWRE PDE ($p = 0.04$). However, none of the indirect mediation effects through either DSF or DSB on the relationship of rise time and reading were significant. In addition, the SEM model with RAN (rapid picture naming) as the mediator, showed a poor fit (*chi-square* $p < 0.01$), suggesting that path coefficients may not accurately reflect the theoretical relationship and should be interpreted with caution. In this model, rise time did not directly affect RAN, although RAN directly impacted various reading measures, including WJ word ID ($p < 0.01$), TOWRE SWE ($p = 0.03$) and TOWRE PDE ($p = 0.02$), with no significant indirect mediation effects observed.

For the relationship between speech in noise and reading, the SEM model with phonological memory, namely DSF and DSB, as mediators indicated a good fit (*chi-square* $p = 0.28$). Speech in noise did not show direct effects on digit spans, but DSB directly impacted on TOWREs (*p.s.* < 0.05). None of the mediating effects through digit span is significant. In addition, the SEM model with RAN as the mediator, however, showed a poor fit ($p < 0.01$), suggesting that path coefficients should also be treated with caution. Specifically, speech in noise did not directly impact RAN while RAN only marginally impacted TOWRE PDE ($p = 0.08$), with no significant indirect mediating effect detected. Overall, our results suggested that neither digit spans nor RAN mediates the relationship between auditory processing and reading outcomes. For more detailed SEM results, see Supplementary Table 9.

#### ART and SiN tasks tap into different linguistic and cognitive abilities

We then aimed to directly characterize subgroups of the DD cohort with specific difficulties in one of the two tasks. To this end, we subset the entire cohort to two subgroups, with particularly low (bottom 25$^{th}$ percentile) or high (upper 25$^{th}$ percentile) performance on each task. As expected, ART-based subgroups ($n = 21$ and 16 for high and low performance) differed in all measures that were found to be correlated with amplitude rise time discrimination abilities in the entire DD group (Table 1 in the main texts), including WJ word ID, phonological sound awareness, and digit span backward scores (Supplementary Table 7). Likewise, SiN-based subgroups ($n = 18$ and 17 for high and low performance) differed in tasks that were correlated with speech in noise abilities in the entire DD cohort, including ROWPVT vocabulary and rapid picture naming, with marginal differences in phonological sound awareness and digit span forward (Supplementary Table 7). Next, direct comparisons of correlations across tasks showed significantly different correlations for digit span backward ($p = 0.02$, $z = 2.37$) and rapid picture naming ($p = 0.01$, $z = 2.49$), whereas all other correlations did not differ significantly across tasks. Overall, this analysis suggests that speech in noise recognition and amplitude rise time discrimination are associated with different domains of verbal memory. That is, verbal working memory is associated with non-speech auditory processing, whereas processing speed is associated with speech perception abilities. In short, this subgrouping analysis replicates the correlational patterns within the DD group and suggests that the two auditory tasks tap into (partially) distinct neurocognitive abilities although psychometric differences exist.

### Supplementary Figures

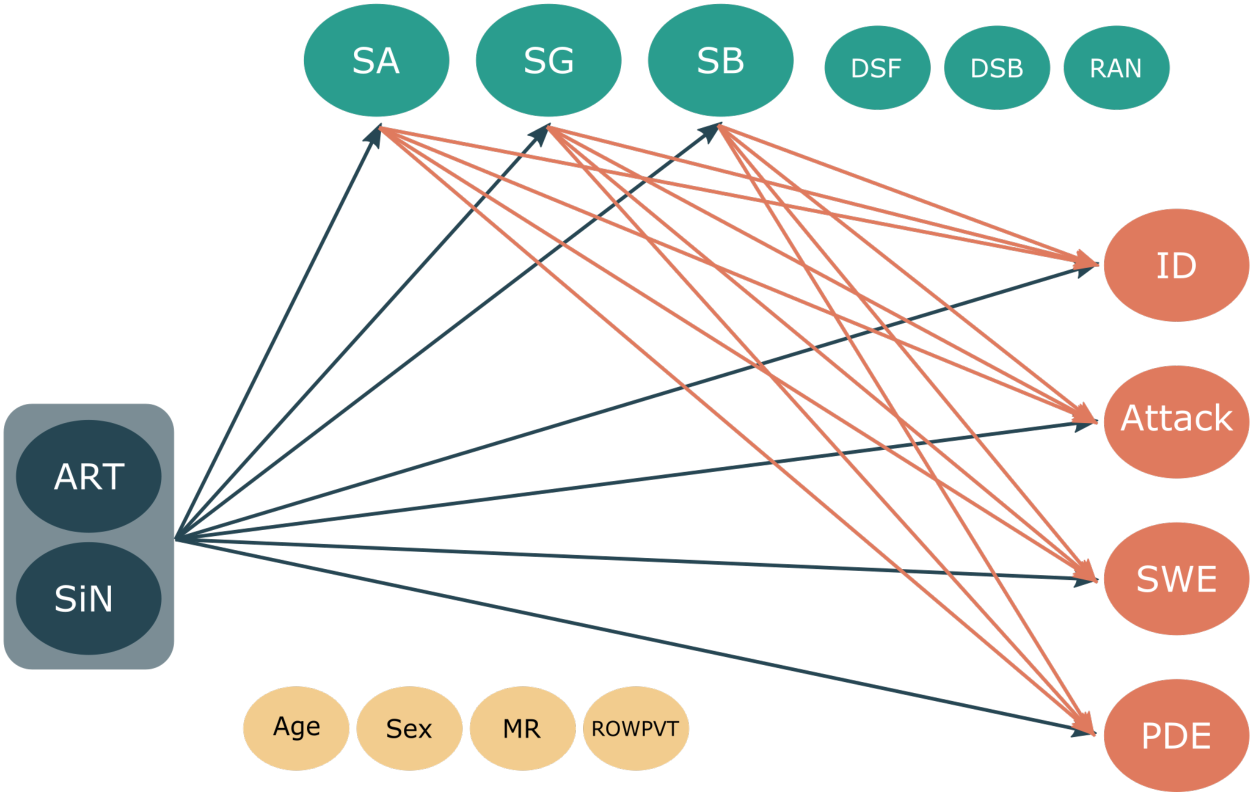


**Supplementary Fig. 1 Schematic of direct acyclic graphs (DAGs) illustrating the causal relationship between auditory abilities and reading outcomes.** That is, amplitude rise time and speech in noise impacts reading through phonology. Dark blue denotes exposures, namely auditory abilities: amplitude rise time (ART) and speech in noise (SiN). Orange denotes reading outcomes, namely: WJ word identification (ID), WJ Attack, TOWRE sight word efficiency (SWE), and TOWRE phonemic decoding efficiency (PDE). Green denotes phonological mediators, including phonological awareness (segmentation, SG; sound awareness, SA; sound blending, SB), digit spans as a proxy of phonological memory, and rapid picture naming (RAN) as a proxy of rapid automatized naming. Yellow denotes covariates of no interest, including age, sex, matrix reasoning (MR), and receptive one-word picture vocabulary (ROWPVT). This DAG was refined by adding direct paths between the four reading outcomes (dotted lines). Of note, due to sample size constraints and to ensure robust estimation, only phonological awareness measures were taken as mediators in the main text. For details of other phonological measures see Supplementary Results and Supplementary Table 9. DSF: digit span forward; DSB: digit span backward.

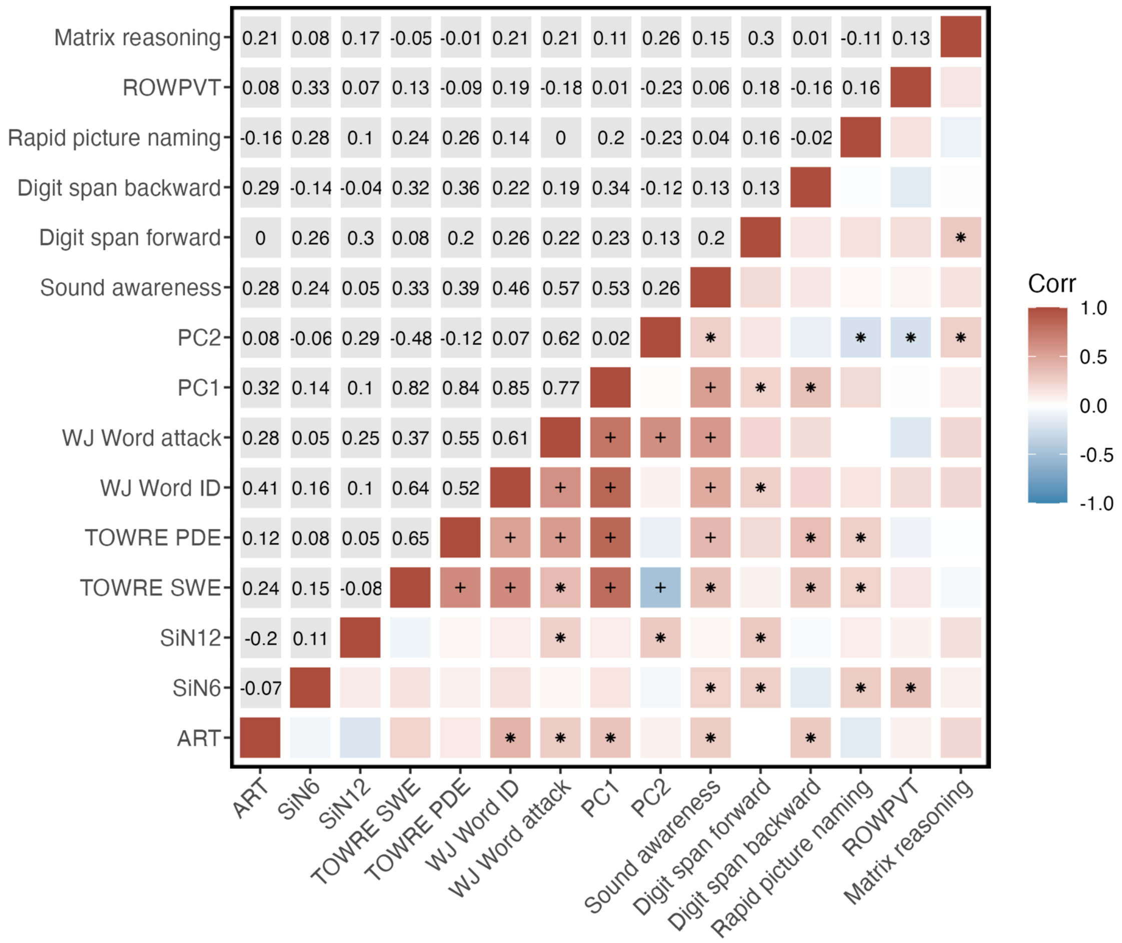


**Supplementary Fig. 2 Correlation matrix between amplitude rise time, speech in noise, and reading (and reading PCs), phonology, and cognitive measures.** The upper panel illustrates the correlation coefficients after controlling for age and sex, and the lower panel illustrates the pattern of correlations. ART: Amplitude rise time, SiN: speech in noise, ROWPVT: receptive one-word picture vocabulary test, TOWRE SWE: single-word reading efficiency, TOWRE PDE: phonemic decoding efficiency, PC1: reading PC1, PC2: reading PC2. + $P < 0.05$ after multiple comparisons correction, * $P < 0.05$.

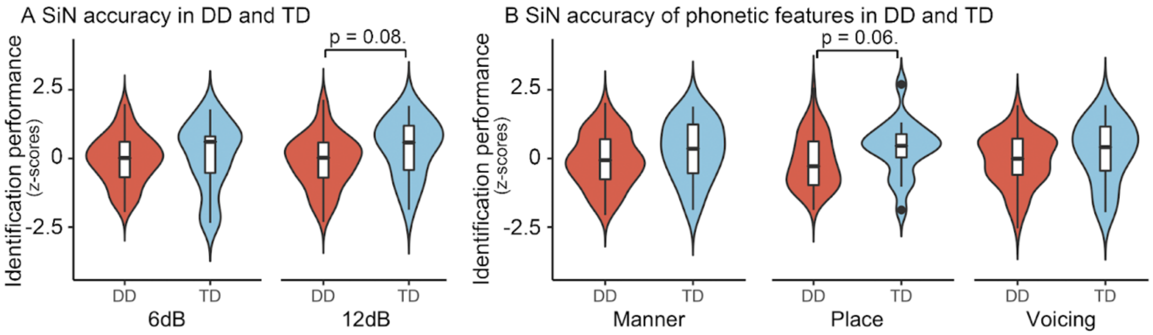


**Supplementary Fig. 3 Speech in noise recognition abilities at 12 dB between DD and TD children.** (**A**) Marginal group differences in speech in noise recognition abilities between DD and TD groups in the 12 dB condition. (**B**) Significant main group and feature effects in the recognition for single phonetic features. Post-hoc analysis revealed selective impairments in DD for the recognition of the place of articulation. The auditory processing ability scores were z-transformed. * denotes $P < 0.05$ and . denotes $P < 0.10$. Red and blue denote the DD and TD, respectively.

### Supplementary Tables

**Supplementary Table 1. The number of children who completed each auditory task and with MRI scans.**

| n | Tasks/MRI scans | DD+TD | DD | TD |
|---|---|---|---|---|
| **Initial sample:** | | | | |
| | Auditory tasks | 135 | 90 | 45 |
| | MRI scans | 126 | 88 | 38 |
| **After screening:** | | | | |
| Matrix reasoning | Auditory tasks | 110 | 78 | 32 |
| | MRI scans | 102 | 76 | 26 |
| ART task[a] | Auditory tasks | 86 | 58 | 28 |
| | MRI scans | 78 | 56 | 22 |
| SiN task at 6 dB[b] | Auditory tasks | 92 | 66 | 26 |
| | MRI scans | 84 | 64 | 20 |
| SiN task at 12dB[b] | Auditory tasks | 81 | 63 | 18 |
| | MRI scans | 75 | 61 | 14 |

[a]For the ART task, children with an accuracy below 65% were excluded from the analysis.

[b]For the SiN task, children who fall beyond the 1.5 IQR were excluded from the analysis.

**Supplementary Table 2. GORT reading scores in developmental dyslexia (DD) and typically developing children (TD).**

| **GORT reading (%ile) [DD/TD][a]** | **DD** | | **TD** | | | |
|---|---|---|---|---|---|---|
| | **Mean ± SD** | **Range** | **Mean ± SD** | **Range** | ***p-values*** | ***t-values*** |
| Rate [78/22] | 16.64 ± 14.80 | 0.40 - 63.00 | 75.84 ± 17.22 | 37.00 - 99.50 | 0.15 | 1.48 |
| Accuracy [78/22] | 11.04 ± 11.78 | 0.04 - 50.00 | 51.98 ± 23.28 | 16.00 - 99.50 | 0.15 | 1.48 |
| Fluency [78/22] | 12.75 ± 12.13 | 0.40 - 50.00 | 65.11 ± 15.64 | 37.00 - 99.50 | 0.13 | 1.56 |
| Comprehension [78/22] | 22.69 ± 18.52 | 0.50 - 75.00 | 55.91 ± 17.49 | 16.00 - 98.00 | 0.20 | 1.30 |

All scores reflect percentiles (%ile) relative to age-matched population data. Welch's two-sample t-tests were conducted to examine the differences between the DD and TD groups for the GORT reading scores.

[a]Numbers in square brackets denote the sample size for each group.

**Supplementary Table 3. Summary of hierarchical regression analysis for amplitude rise time (ART) predicting reading and phonology in children with DD.**

| DVs | Predictors | β | t | Pr (>\|t\|) | Model F (p) | Adjusted $R^2$ | Adjusted $\Delta R^2$ |
|---|---|---|---|---|---|---|---|
| **WJ word ID** | | | | | | | |
| Step1 | Sex | -0.02 | -0.11 | 0.91 | 1.53 (0.21) | 0.03 | |
| | Age | 0.27 | 1.78 | 0.08 | | | |
| | Matrix reasoning | 0.24 | 1.53 | 0.13 | | | |
| | ROWPVT vocabulary | 0.16 | 1.17 | 0.25 | | | |
| Step2 | ART (z-scored) | 0.35 | 2.74 | <0.01* | 4.27 (<0.01) | 0.15 | 0.11 |
| **WJ word attack** | | | | | | | |
| Step1 | Sex | 0.01 | 0.08 | 0.93 | | | |
| | Age | 0.09 | 0.62 | 0.54 | | | |
| | Matrix reasoning | 0.31 | 1.93 | 0.06. | | | |
| | ROWPVT vocabulary | -0.22 | -1.56 | 0.12 | 1.36 (0.26) | 0.03 | |
| Step2 | ART (z-scored) | 0.24 | 1.75 | 0.09. | 1.75 (0.14) | 0.06 | 0.04 |
| **TOWRE SWE** | | | | | | | |
| Step1 | Sex | 0.05 | 0.33 | 0.74 | | | |
| | Age | 0.25 | 1.65 | 0.10 | | | |
| | Matrix reasoning | -0.02 | -0.10 | 0.92 | | | |
| | ROWPVT vocabulary | 0.11 | 0.78 | 0.43 | 1.06 (0.39) | 0.00 | |
| Step2 | ART (z-scored) | 0.24 | 1.73 | 0.09. | 1.48 (0.21) | 0.04 | 0.04 |
| **TOWRE PDE** | | | | | | | |
| Step1 | Sex | 0.09 | 0.65 | 0.52 | | | |
| | Age | 0.17 | 1.12 | 0.27 | | | |
| | Matrix reasoning | 0.13 | 0.83 | 0.41 | | | |
| | ROWPVT vocabulary | -0.16 | -1.10 | 0.28 | 0.87 (0.49) | -0.01 | |
| Step2 | ART (z-scored) | 0.11 | 0.78 | 0.44 | 0.81 (0.55) | -0.02 | -0.01 |
| **Sound Awareness** | | | | | | | |
| Step1 | Sex | 0.03 | 0.22 | 0.83 | | | |
| | Age | -0.06 | -0.42 | 0.68 | | | |
| | Matrix reasoning | 0.13 | 0.81 | 0.42 | | | |
| | ROWPVT vocabulary | 0.07 | 0.49 | 0.63 | 0.57 (0.69) | -0.03 | |
| Step2 | ART (z-scored) | 0.26 | 1.87 | 0.07. | 1.18 (0.33) | 0.02 | 0.05 |
| **Sound Blending** | | | | | | | |
| Step1 | Sex | 0.01 | 0.05 | 0.97 | | | |
| | Age | -0.27 | -1.92 | 0.06. | | | |
| | Matrix reasoning | 0.15 | 0.99 | 0.33 | | | |
| | ROWPVT vocabulary | 0.22 | 1.69 | 0.10. | 3.41 (0.02) | 0.15 | |
| Step2 | ART (z-scored) | 0.30 | 2.43 | 0.02* | 4.17 (<0.01) | 0.22 | 0.07 |

| | | | | | | | |
|---|---|---|---|---|---|---|---|
| **Segmentation** | | | | | | | |
| Step1 | Sex | -0.14 | -1.11 | 0.27 | | | |
| | Age | -0.20 | -1.46 | 0.15 | | | |
| | Matrix reasoning | 0.43 | 3.04 | <0.01* | | | |
| | ROWPVT vocabulary | -0.05 | -0.41 | 0.69 | 5.02 (<0.01) | 0.23 | |
| Step2 | ART (z-scored) | 0.38 | 3.39 | <0.01* | 7.15 (<0.01) | 0.36 | 0.13 |
| **Rapid picture naming** | | | | | | | |
| Step1 | Sex | -0.15 | -1.09 | 0.28 | | | |
| | Age | -0.35 | -2.42 | 0.02* | | | |
| | Matrix reasoning | -0.21 | -1.41 | 0.17 | | | |
| | ROWPVT vocabulary | 0.15 | 1.09 | 0.28 | 2.56 (<0.05) | 0.10 | |
| Step2 | ART (z-scored) | -0.13 | -0.99 | 0.33 | 2.24 (0.06) | 0.10 | 0.00 |
| **Digit span forward** | | | | | | | |
| Step1 | Sex | 0.07 | 0.51 | 0.62 | | | |
| | Age | -0.14 | -1.00 | 0.32 | | | |
| | Matrix reasoning | 0.36 | 2.48 | 0.02* | | | |
| | ROWPVT vocabulary | 0.10 | 0.81 | 0.42 | 4.05 (<0.01) | 0.18 | |
| Step2 | ART (z-scored) | -0.09 | -0.68 | 0.50 | 3.30 (0.01) | 0.17 | -0.01 |
| **Digit span backward** | | | | | | | |
| Step1 | Sex | 0.24 | 1.70 | 0.10. | | | |
| | Age | 0.12 | 0.81 | 0.42 | | | |
| | Matrix reasoning | 0.09 | 0.59 | 0.56 | | | |
| | ROWPVT vocabulary | -0.09 | -0.63 | 0.53 | 1.29 (0.29) | 0.02 | |
| Step2 | ART (z-scored) | 0.26 | 1.97 | 0.06. | 1.86 (0.12) | 0.07 | 0.05 |

* *P* < 0.05 and . *P* < 0.10.

**Supplementary Table 4. SEM Overview: amplitude rise time (ART) contributes to reading via phonological awareness.**

| Path | Regressor | Outcome | Estimate | Std.Err | z-value | P(>\|z\|) |
|---|---|---|---|---|---|---|
| **Direct effects (A)** | ART | SG | 0.38 | 0.11 | 3.54 | <0.01* |
| | | SB | 0.30 | 0.13 | 2.31 | 0.02* |
| | | SA | 0.26 | 0.16 | 1.61 | 0.11 |
| **Direct effects (B)** | SG | WJ ID | -0.09 | 0.14 | -0.63 | 0.53 |
| | | WJ Attack | 0.06 | 0.13 | 0.43 | 0.67 |
| | | TOWRE SWE | -0.38 | 0.16 | -2.30 | 0.02* |
| | | TOWRE PDE | -0.10 | 0.18 | -0.57 | 0.57 |
| | SA | WJ ID | 0.38 | 0.12 | 3.14 | <0.01* |
| | | WJ Attack | 0.53 | 0.10 | 5.52 | <0.01* |
| | | TOWRE SWE | 0.37 | 0.14 | 2.74 | <0.01* |
| | | TOWRE PDE | 0.40 | 0.12 | 3.46 | <0.01* |
| | SB | WJ ID | 0.15 | 0.13 | 1.16 | 0.25 |
| | | WJ Attack | 0.25 | 0.12 | 2.10 | 0.04* |
| | | TOWRE SWE | -0.06 | 0.16 | -0.37 | 0.72 |
| | | TOWRE PDE | 0.06 | 0.11 | 0.52 | 0.61 |
| **Direct effects (C')** | ART | WJ ID | 0.25 | 0.13 | 1.94 | 0.05. |
| | | WJ Attack | 0.00 | 0.11 | 0.03 | 0.98 |
| | | TOWRE SWE | 0.30 | 0.15 | 2.06 | 0.04* |
| | | TOWRE PDE | 0.03 | 0.14 | 0.20 | 0.84 |
| **Indirect effects** | ART -> SG | WJ ID | -0.03 | 0.06 | -0.58 | 0.56 |
| | ART -> SB | WJ ID | 0.04 | 0.04 | 1.07 | 0.28 |
| | ART -> SA | WJ ID | 0.10 | 0.08 | 1.29 | 0.20 |
| | ART -> SG | WJ Attack | 0.02 | 0.05 | 0.39 | 0.70 |
| | ART -> SB | WJ Attack | 0.08 | 0.04 | 1.77 | 0.08. |
| | ART -> SA | WJ Attack | 0.14 | 0.09 | 1.56 | 0.12 |
| | ART -> SG | TOWRE SWE | -0.14 | 0.07 | -1.99 | <0.05* |
| | ART -> SB | TOWRE SWE | -0.02 | 0.05 | -0.33 | 0.75 |
| | ART -> SA | TOWRE SWE | 0.10 | 0.08 | 1.25 | 0.21 |
| | ART -> SG | TOWRE PDE | -0.04 | 0.07 | -0.56 | 0.58 |
| | ART -> SB | TOWRE PDE | 0.02 | 0.04 | 0.50 | 0.62 |
| | ART -> SA | TOWRE PDE | 0.10 | 0.07 | 1.47 | 0.14 |
| **Total effects (C)** | ART | WJ ID | 0.35 | 0.13 | 2.81 | <0.01* |
| | | WJ Attack | 0.24 | 0.15 | 1.56 | 0.12 |
| | | TOWRE SWE | 0.24 | 0.14 | 1.72 | 0.09. |
| | | TOWRE PDE | 0.11 | 0.15 | 0.75 | 0.45 |

A paths: direct effects of ART on mediators (sound awareness [SA], blending [SB], segmentation [SG]); B paths: mediators' impact on reading; C' paths: direct ART effects on reading, controlling for mediators; C paths: total effects of ART on reading; Indirect effects: product of A and B paths, indicating mediation. WJ ID: WJ word identification; WJ Attack: WJ word attack; TOWRE SWE: TOWRE sight word efficiency; TOWRE PDE: TOWRE phonemic decoding efficiency. * $P < 0.05$, . $P < 0.10$.

**Supplementary Table 5. Summary of hierarchical regression analysis for speech in noise (SiN) predicting reading and phonology in children with DD.**

| DVs | Predictors | β | t | Pr (>\|t\|) | Model F (p) | Adjusted R2 | Adjusted ΔR2 |
|---|---|---|---|---|---|---|---|
| **WJ word ID** | | | | | | | |
| Step1 | Sex | 0.02 | 0.16 | 0.87 | | | |
| | Age | 0.36 | 2.64 | 0.01* | | | |
| | Matrix reasoning | 0.25 | 0.17 | 0.09. | | | |
| | ROWPVT vocabulary | 0.16 | 1.26 | 0.21 | 2.70 (0.04) | 0.10 | |
| Step2 | SiN (z-scored) | 0.10 | 0.71 | 0.48 | 2.24 (0.06) | 0.09 | -0.01 |
| **WJ word attack** | | | | | | | |
| Step1 | Sex | 0.10 | 0.78 | 0.44 | | | |
| | Age | 0.21 | 1.54 | 0.13 | | | |
| | Matrix reasoning | 0.29 | 2.04 | <0.05* | | | |
| | ROWPVT vocabulary | -0.23 | -1.85 | 0.07. | 2.84 (0.03) | 0.11 | |
| Step2 | SiN (z-scored) | 0.12 | 0.90 | 0.37 | 2.43 (<0.05) | 0.10 | 0.00 |
| **TOWRE SWE** | | | | | | | |
| Step1 | Sex | 0.10 | -0.55 | 0.49 | | | |
| | Age | 0.31 | 0.70 | 0.03* | | | |
| | Matrix reasoning | -0.11 | 2.20 | 0.47 | | | |
| | ROWPVT vocabulary | 0.11 | -0.74 | 0.38 | 2.35 (0.06) | 0.08 | |
| Step2 | SiN (z-scored) | 0.14 | 0.99 | 0.33 | 2.08 (0.08) | 0.09 | 0.01 |
| **TOWRE PDE** | | | | | | | |
| Step1 | Sex | 0.15 | 1.05 | 0.30 | | | |
| | Age | 0.22 | 1.57 | 0.12 | | | |
| | Matrix reasoning | 0.02 | 0.10 | 0.92 | | | |
| | ROWPVT vocabulary | -0.12 | -0.96 | 0.34 | 1.62 (0.18) | 0.04 | |
| Step2 | SiN (z-scored) | 0.14 | 0.95 | 0.34 | 1.48 (0.21) | 0.04 | 0.00 |
| **Sound Awareness** | | | | | | | |
| Step1 | Sex | 0.16 | 1.13 | 0.26 | | | |
| | Age | 0.07 | 0.46 | 0.65 | | | |
| | Matrix reasoning | 0.21 | 1.37 | 0.18 | | | |
| | ROWPVT vocabulary | 0.00 | -0.01 | 0.99 | 1.37 (0.26) | 0.02 | |
| Step2 | SiN (z-scored) | 0.29 | 2.09 | 0.04* | 2.03 (0.09) | 0.08 | 0.05 |
| **Sound Blending** | | | | | | | |
| Step1 | Sex | 0.06 | 0.49 | 0.63 | | | |
| | Age | -0.08 | -0.58 | 0.57 | | | |
| | Matrix reasoning | 0.31 | 2.21 | 0.03* | | | |

| | | | | | | | |
|---|---|---|---|---|---|---|---|
| | ROWPVT vocabulary | 0.23 | 1.90 | 0.06. | 3.86 (<0.01) | 0.16 | |
| Step2 | SiN (z-scored) | -0.02 | -0.16 | 0.88 | 3.04 (0.02) | 0.14 | -0.01 |
| **Segmentation** | | | | | | | |
| Step1 | Sex | -0.13 | -0.96 | 0.34 | | | |
| | Age | -0.11 | -0.84 | 0.40 | | | |
| | Matrix reasoning | 0.39 | 2.72 | <0.01* | | | |
| | ROWPVT vocabulary | -0.02 | -0.13 | 0.90 | 3.19 (0.02) | 0.13 | |
| Step2 | SiN (z-scored) | -0.07 | -0.54 | 0.59 | 2.58 (0.19) | 0.11 | -0.01 |
| **Rapid picture naming** | | | | | | | |
| Step1 | Sex | -0.14 | -1.03 | 0.31 | | | |
| | Age | -0.11 | -0.79 | 0.44 | | | |
| | Matrix reasoning | -0.12 | -0.80 | 0.43 | | | |
| | ROWPVT vocabulary | 0.24 | 1.89 | 0.06. | 1.75 (0.15) | 0.05 | |
| Step2 | SiN (z-scored) | 0.23 | 1.65 | 0.11 | 1.98 (<0.10) | 0.07 | 0.03 |
| **Digit span forward** | | | | | | | |
| Step1 | Sex | 0.04 | 0.33 | 0.74 | | | |
| | Age | -0.02 | -0.11 | 0.91 | | | |
| | Matrix reasoning | 0.34 | 2.37 | 0.02* | | | |
| | ROWPVT vocabulary | 0.22 | 1.81 | 0.08. | 3.55 (0.012) | 0.14 | |
| Step2 | SiN (z-scored) | 0.21 | 1.57 | 0.12 | 3.41 (<0.01) | 0.16 | 0.02 |
| **Digit span backward** | | | | | | | |
| Step1 | Sex | 0.31 | 2.19- | 0.03* | | | |
| | Age | 0.00 | -0.04 | 0.97 | | | |
| | Matrix reasoning | -0.03 | -0.21 | 0.84 | | | |
| | ROWPVT vocabulary | -0.13 | -1.05 | 0.30 | 1.71 (0.16) | 0.04 | |
| Step2 | SiN (z-scored) | -0.10 | -0.67 | 0.51 | 1.44 (0.22) | 0.03 | -0.01 |

* $P < 0.05$ and . $P < 0.10$.

**Supplementary Table 6. SEM Overview: speech in noise (SiN) contributes to reading via phonological awareness.**

| Path | Regressor | Outcome | Estimate | Std.Err | z-value | P(>\|z\|) |
|---|---|---|---|---|---|---|
| **Direct effects (A)** | SiN | SG | -0.07 | 0.15 | -0.50 | 0.62 |
| | | SB | -0.02 | 0.13 | -0.17 | 0.87 |
| | | SA | 0.29 | 0.12 | 2.47 | 0.01* |
| **Direct effects (B)** | SG | WJ ID | -0.05 | 0.13 | -0.36 | 0.73 |
| | | WJ Attack | 0.04 | 0.11 | 0.33 | 0.74 |
| | | TOWRE SWE | -0.20 | 0.15 | -1.36 | 0.17 |
| | | TOWRE PDE | -0.01 | 0.15 | -0.06 | 0.95 |
| | SA | WJ ID | 0.38 | 0.13 | 3.06 | <0.01* |
| | | WJ Attack | 0.50 | 0.10 | 5.09 | <0.01* |
| | | TOWRE SWE | 0.33 | 0.15 | 2.11 | 0.04* |
| | | TOWRE PDE | 0.36 | 0.12 | 3.02 | <0.01* |
| | SB | WJ ID | 0.16 | 0.13 | 1.26 | 0.21 |
| | | WJ Attack | 0.14 | 0.11 | 1.26 | 0.21 |
| | | TOWRE SWE | -0.02 | 0.15 | -0.16 | 0.87 |
| | | TOWRE PDE | -0.04 | 0.11 | -0.32 | 0.75 |
| **Direct effects (C')** | SiN | WJ ID | -0.01 | 0.13 | -0.11 | 0.92 |
| | | WJ Attack | -0.02 | 0.11 | -0.14 | 0.89 |
| | | TOWRE SWE | 0.03 | 0.15 | 0.19 | 0.85 |
| | | TOWRE PDE | 0.03 | 0.11 | 0.26 | 0.79 |
| **Indirect effects** | SiN->SG | WJ ID | 0.00 | 0.03 | 0.14 | 0.89 |
| | SiN->SB | WJ ID | 0.00 | 0.03 | -0.12 | 0.90 |
| | SiN->SA | WJ ID | 0.11 | 0.06 | 1.99 | <0.05* |
| | SiN->SG | WJ Attack | 0.00 | 0.02 | -0.13 | 0.89 |
| | SiN->SB | WJ Attack | 0.00 | 0.02 | -0.13 | 0.90 |
| | SiN->SA | WJ Attack | 0.15 | 0.07 | 2.07 | 0.04* |
| | SiN->SG | TOWRE SWE | 0.02 | 0.04 | 0.39 | 0.70 |
| | SiN->SB | TOWRE SWE | 0.00 | 0.02 | 0.03 | 0.98 |
| | SiN->SA | TOWRE SWE | 0.10 | 0.06 | 1.58 | 0.12 |
| | SiN->SG | TOWRE PDE | 0.00 | 0.03 | 0.03 | 0.98 |
| | SiN->SB | TOWRE PDE | 0.00 | 0.01 | 0.05 | 0.96 |
| | SiN->SA | TOWRE PDE | 0.11 | 0.06 | 1.87 | 0.06. |
| **Total effects (C)** | SiN | WJ ID | 0.10 | 0.13 | 0.79 | 0.43 |
| | | WJ Attack | 0.13 | 0.14 | 0.91 | 0.37 |
| | | TOWRE SWE | 0.10 | 0.14 | 0.71 | 0.48 |
| | | TOWRE PDE | 0.09 | 0.14 | 0.69 | 0.49 |

A paths: the effects of SiN on the mediators (sound awareness [SA], blending [SB], segmentation [SG]); B paths: the effects of mediators on reading; C' paths: the direct effects of SiN on reading; C paths: the total effect of SiN on reading without considering the mediation effect; indirect effects are the product of a and b paths, showing the mediation effects. WJ ID: WJ word identification; WJ Attack: WJ word attack; TOWRE SWE: TOWRE sight word efficiency; TOWRE PDE: TOWRE Phonemic decoding efficiency. * $P < 0.05$, . $P < 0.10$.

**Supplementary Table 7. Group differences between two subgroups of DD formulated according to the performance in amplitude rise time (ART) and speech in noise (SiN) tasks.**

| | **ART DD subgroups** | | | **SiN DD subgroups** | | |
|---|---|---|---|---|---|---|
| | **High (n = 21)** | **Low (n = 16)** | | **High (n = 18)** | **Low (n = 17)** | |
| | Mean±SD | Mean±SD | *p-values* | Mean±SD | Mean±SD | *p-values* |
| **Demographics** | | | | | | |
| Age (years) | 11.27±1.88 | 10.71±1.72 | 0.35 | | 11.82±1.79 | 10.08±1.80 |
| Sex (F/M) | 8/13 | 5/11 | 0.89 | | 8/10 | 6/11 |
| **Experimental Auditory task** | | | | | | |
| ART | 65.95±27.17 | 437.42±51.72 | <0.01* | 254.24±155.01 | 256.17±173.72 | 0.63 |
| SiN (%) | 42.59±8.27 | 42.77±5.61 | 0.74 | 51.85±3.79 | 32.98±3.39 | <0.01* |
| **Single word/nonword reading (%ile)** | | | | | | |
| *TOWRE-2* | | | | | | |
| TOWRE SWE | 24.46±22.94 | 11.26±12.68 | 0.06. | 23.32±22.30 | 11.68±18.37 | 0.39 |
| TOWRE PDE | 22.22±21.85 | 15.06±17.46 | 0.32 | 17.94±17.27 | 13.41±20.60 | 0.89 |
| *WJ-IV* | | | | | | |
| Word ID | 40.20±27.46 | 17.41±16.42 | <0.01* | 31.64±27.60 | 20.65±23.92 | 0.48 |
| Word attack | 49.76±23.74 | 33.63±25.23 | 0.06. | 36.94±27.58 | 37.00±23.96 | 0.84 |
| **Phonological awareness (%ile)** | | | | | | |
| Segmentation | 70.43±19.26 | 51.00±22.37 | <0.01* | 51.67±23.88 | 62.59±19.65 | 0.39 |
| Sound blending | 69.00±22.17 | 50.38±22.74 | <0.01* | 49.17±24.93 | 53.76±28.56 | 0.96 |
| Sound awareness | 48.57±26.82 | 30.75±26.09 | 0.04* | 45.39±28.97 | 30.24±26.01 | 0.09. |
| **Cognitive and language (%ile)** | | | | | | |
| Matrix reasoning | 70.67±21.76 | 63.56±19.52 | 0.07. | 64.06±23.51 | 68.47±22.74 | 0.32 |
| Digit span forward | 32.90±28.60 | 31.38±21.57 | 0.59 | 34.61±27.27 | 24.47±21.08 | 0.08. |
| Digit span backward | 47.10±24.30 | 31.25±19.88 | 0.03* | 35.22±23.00 | 42.24±27.49 | 0.45 |
| ROWPVT vocabulary | 69.40±24.57 | 66.45±24.98 | 0.66 | 72.72±22.68 | 58.04±25.63 | 0.05* |
| Rapid picture naming | 25.62±22.44 | 34.38±22.04 | 0.28 | 33.83±17.86 | 20.24±22.82 | 0.03* |

Welch's two-sample t-test was used to compare two DD subgroups. Fisher's exact test was conducted for the group difference regarding sex. Age and sex were controlled for in the group comparison. All reading, language, and cognitive scores reflect percentiles (%ile) relative to age-matched population data.

* $P < 0.05$ and . $P < 0.10$.

**Supplementary Table 8. SEM Overview: amplitude rise time (ART) and speech in noise (SiN) contributes to reading via phonological awareness.**

| Path | Regressor | Outcome | Estimate | Std.Err | z-value | P(>\|z\|) |
|---|---|---|---|---|---|---|
| **Direct effects (A)** | ART | SG | 0.36 | 0.11 | 3.47 | <0.01* |
| | | SB | 0.34 | 0.14 | 2.54 | 0.01* |
| | | SA | 0.20 | 0.17 | 1.17 | 0.24 |
| | SiN | SG | -0.08 | 0.14 | -0.54 | 0.59 |
| | | SB | 0.03 | 0.13 | 0.22 | 0.83 |
| | | SA | 0.35 | 0.14 | 2.48 | 0.01* |
| **Direct effects (B)** | SG | WJ ID | -0.07 | 0.16 | -0.44 | 0.66 |
| | | WJ Attack | 0.05 | 0.14 | 0.37 | 0.71 |
| | | TOWRE SWE | -0.39 | 0.18 | -2.21 | 0.03* |
| | | TOWRE PDE | -0.11 | 0.19 | -0.58 | 0.56 |
| | SA | WJ ID | 0.41 | 0.13 | 3.23 | <0.01* |
| | | WJ Attack | 0.53 | 0.11 | 4.88 | <0.01* |
| | | TOWRE SWE | 0.35 | 0.16 | 2.18 | 0.03* |
| | | TOWRE PDE | 0.37 | 0.13 | 2.80 | <0.01* |
| | SB | WJ ID | 0.08 | 0.14 | 0.55 | 0.58 |
| | | WJ Attack | 0.23 | 0.13 | 1.76 | 0.08. |
| | | TOWRE SWE | -0.10 | 0.17 | -0.59 | 0.55 |
| | | TOWRE PDE | 0.04 | 0.12 | 0.30 | 0.76 |
| **Direct effects (C')** | ART | WJ ID | 0.27 | 0.14 | 1.93 | 0.05. |
| | | WJ Attack | -0.01 | 0.12 | -0.07 | 0.95 |
| | | TOWRE SWE | 0.32 | 0.16 | 2.04 | 0.04* |
| | | TOWRE PDE | 0.03 | 0.15 | 0.17 | 0.87 |
| | SiN | WJ ID | -0.07 | 0.14 | -0.50 | 0.62 |
| | | WJ Attack | -0.09 | 0.13 | -0.68 | 0.50 |
| | | TOWRE SWE | -0.03 | 0.16 | -0.20 | 0.84 |
| | | TOWRE PDE | -0.04 | 0.12 | -0.35 | 0.73 |
| **Indirect effects** | ART->SG | WJ ID | -0.03 | 0.06 | -0.41 | 0.68 |
| | ART->SB | WJ ID | 0.03 | 0.05 | 0.53 | 0.60 |
| | ART->SA | WJ ID | 0.08 | 0.08 | 1.02 | 0.31 |
| | ART->SG | WJ Attack | 0.02 | 0.06 | 0.34 | 0.73 |
| | ART->SB | WJ Attack | 0.08 | 0.05 | 1.52 | 0.13 |
| | ART->SA | WJ Attack | 0.11 | 0.09 | 1.15 | 0.25 |
| | ART->SG | TOWRE SWE | -0.14 | 0.07 | -1.98 | <0.05* |
| | ART->SB | TOWRE SWE | -0.03 | 0.06 | -0.53 | 0.59 |
| | ART->SA | TOWRE SWE | 0.07 | 0.08 | 0.92 | 0.36 |

| | | | | | | |
|---|---|---|---|---|---|---|
| | ART->SG | TOWRE PDE | -0.04 | 0.07 | -0.57 | 0.57 |
| | ART->SB | TOWRE PDE | 0.01 | 0.04 | 0.29 | 0.77 |
| | ART->SA | TOWRE PDE | 0.07 | 0.07 | 1.08 | 0.28 |
| **Indirect effects** | SiN->SG | WJ ID | 0.01 | 0.03 | 0.17 | 0.86 |
| | SiN->SB | WJ ID | 0.00 | 0.02 | 0.10 | 0.92 |
| | SiN->SA | WJ ID | 0.14 | 0.07 | 2.15 | 0.03* |
| | SiN->SG | WJ Attack | 0.00 | 0.03 | -0.16 | 0.87 |
| | SiN->SB | WJ Attack | 0.01 | 0.03 | 0.19 | 0.85 |
| | SiN->SA | WJ Attack | 0.18 | 0.09 | 2.07 | 0.04* |
| | SiN->SG | TOWRE SWE | 0.03 | 0.06 | 0.48 | 0.63 |
| | SiN->SB | TOWRE SWE | 0.00 | 0.03 | -0.11 | 0.91 |
| | SiN->SA | TOWRE SWE | 0.12 | 0.08 | 1.51 | 0.13 |
| | SiN->SG | TOWRE PDE | 0.01 | 0.04 | 0.24 | 0.81 |
| | SiN->SB | TOWRE PDE | 0.00 | 0.02 | 0.06 | 0.95 |
| | SiN->SA | TOWRE PDE | 0.13 | 0.07 | 1.79 | 0.07. |
| **Total effects (C)** | ART | WJ ID | 0.35 | 0.14 | 2.56 | 0.01* |
| | | WJ Attack | 0.20 | 0.16 | 1.22 | 0.22 |
| | | TOWRE SWE | 0.22 | 0.14 | 1.52 | 0.13 |
| | | TOWRE PDE | 0.07 | 0.15 | 0.46 | 0.64 |
| **Total effects (C)** | SiN | WJ ID | 0.08 | 0.15 | 0.52 | 0.60 |
| | | WJ Attack | 0.10 | 0.13 | 0.78 | 0.43 |
| | | TOWRE SWE | 0.12 | 0.15 | 0.78 | 0.44 |
| | | TOWRE PDE | 0.09 | 0.13 | 0.73 | 0.46 |

A paths represent the effects of two auditory processing abilities on the mediators (sound awareness [SA], blending [SB], segmentation [SG]); B paths indicate the effects of mediators on reading; C' paths represent the direct effects of auditory processing abilities on reading; C paths indicate the total effect of auditory processing abilities on reading without considering the mediation effect; indirect effects are the product of a and b paths, showing the mediation effects. WJ ID: WJ word identification; WJ Attack: WJ word attack; TOWRE SWE: TOWRE sight word efficiency; TOWRE PDE: TOWRE Phonemic decoding efficiency. * $P < 0.05$, . $P < 0.10$.

**Supplementary Table 9. SEM Overview: amplitude rise time (ART) and speech in noise (SiN) contributes to reading via phonological memory and rapid automatized naming, respectively.**

| Path | Outcome | Regressor | Estimate | Std.Err | z-value | P(>\|z\|) | Regressor | Estimate | Std.Err | z-value | P(>\|z\|) |
|---|---|---|---|---|---|---|---|---|---|---|---|
| **Direct effects (A)** | RAN | ART | -0.13 | 0.13 | -1.02 | 0.31 | SiN | 0.23 | 0.15 | 1.55 | 0.12 |
| | DSF | | -0.09 | 0.12 | -0.74 | 0.46 | | 0.21 | 0.12 | 1.70 | 0.09. |
| | DSB | | 0.26 | 0.13 | 2.09 | 0.04* | | -0.10 | 0.14 | -0.68 | 0.50 |
| **Direct effects (B)** | WJ ID | RAN | 0.38 | 0.13 | 2.98 | <0.01* | RAN | 0.18 | 0.17 | 1.08 | 0.28 |
| | WJ Attack | | 0.17 | 0.12 | 1.46 | 0.15 | | 0.10 | 0.13 | 0.79 | 0.43 |
| | TOWRE SWE | | 0.33 | 0.15 | 2.20 | 0.03* | | 0.24 | 0.15 | 1.58 | 0.11 |
| | TOWRE PDE | | 0.33 | 0.14 | 2.43 | 0.02* | | 0.25 | 0.14 | 1.78 | 0.08. |
| | WJ ID | DSF | 0.17 | 0.16 | 1.07 | 0.29 | DSF | 0.09 | 0.14 | 0.63 | 0.53 |
| | WJ Attack | | 0.15 | 0.14 | 1.12 | 0.26 | | 0.08 | 0.12 | 0.65 | 0.52 |
| | TOWRE SWE | | -0.02 | 0.18 | -0.10 | 0.92 | | -0.12 | 0.16 | -0.73 | 0.47 |
| | TOWRE PDE | | 0.19 | 0.15 | 1.30 | 0.20 | | 0.10 | 0.14 | 0.72 | 0.47 |
| | WJ ID | DSB | 0.13 | 0.15 | 0.90 | 0.37 | DSB | 0.20 | 0.12 | 1.63 | 0.10 |
| | WJ Attack | | 0.05 | 0.14 | 0.35 | 0.73 | | 0.11 | 0.11 | 0.95 | 0.34 |
| | TOWRE SWE | | 0.24 | 0.13 | 1.77 | 0.08. | | 0.32 | 0.12 | 2.57 | 0.01* |
| | TOWRE PDE | | 0.28 | 0.14 | 2.03 | 0.04* | | 0.30 | 0.13 | 2.33 | 0.02* |
| **Indirect effects** | WJ ID | ART->RAN | -0.05 | 0.05 | -0.94 | 0.35 | SiN->RAN | 0.04 | 0.05 | 0.90 | 0.37 |
| | WJ Attack | ART->RAN | -0.023 | 0.03 | -0.77 | 0.44 | SiN->RAN | 0.02 | 0.04 | 0.62 | 0.53 |
| | TOWRE SWE | ART->RAN | -0.043 | 0.05 | -0.85 | 0.40 | SiN->RAN | 0.06 | 0.05 | 1.04 | 0.30 |
| | TOWRE PDE | ART->RAN | -0.043 | 0.05 | -0.86 | 0.39 | SiN->RAN | 0.06 | 0.06 | 1.02 | 0.31 |
| | WJ ID | ART->DSF | -0.02 | 0.03 | -0.43 | 0.67 | SiN->DSF | 0.02 | 0.04 | 0.55 | 0.58 |
| | WJ ID | ART->DSB | 0.06 | 0.07 | 0.89 | 0.37 | SiN->DSB | -0.01 | 0.03 | -0.43 | 0.67 |
| | WJ Attack | ART->DSF | -0.01 | 0.03 | -0.45 | 0.65 | SiN->DSF | 0.02 | 0.03 | 0.54 | 0.59 |
| | WJ Attack | ART->DSB | 0.01 | 0.04 | 0.32 | 0.75 | SiN->DSB | -0.01 | 0.02 | -0.44 | 0.66 |
| | TOWRE SWE | ART->DSF | 0.00 | 0.03 | 0.06 | 0.96 | SiN->DSF | -0.02 | 0.04 | -0.63 | 0.53 |
| | TOWRE SWE | ART->DSB | 0.06 | 0.05 | 1.34 | 0.18 | SiN->DSB | -0.03 | 0.05 | -0.63 | 0.53 |
| | TOWRE PDE | ART->DSF | -0.02 | 0.03 | -0.50 | 0.62 | SiN->DSF | 0.02 | 0.03 | 0.62 | 0.54 |
| | TOWRE PDE | ART->DSB | 0.07 | 0.05 | 1.45 | 0.15 | SiN->DSB | -0.03 | 0.05 | -0.62 | 0.54 |

Of note, for each auditory processing ability, we performed separate SEM analyses of the mediating effects of phonological memory (digit span backward [DSB]; DSF: digit span forward [DSF]) and rapid automatized naming (rapid picture naming [RAN]) on the relationship between auditory processing abilities and reading. For better presentation, we integrated the results into one table. See Table 2 in the main text for the direct effects and total effects of each auditory processing ability on reading.

A paths: the effects of auditory processing abilities on the mediators (digit spans and rapid picture naming); indirect effects are the product of a and b paths, showing the mediation effects. WJ ID: WJ word identification; WJ Attack: WJ word attack; TOWRE SWE: TOWRE sight word efficiency; TOWRE PDE: TOWRE Phonemic decoding efficiency. * $P < 0.05$, . $P < 0.10$.
